# Quantifying the evolutionary paths to endomembranes

**DOI:** 10.1101/2024.04.15.589612

**Authors:** Paul E. Schavemaker, Michael Lynch

**Affiliations:** Biodesign Center for Mechanisms of Evolution, Arizona State University, Tempe, AZ, 85287, USA

## Abstract

Eukaryotes exhibit a complex and dynamic internal meshwork of membranes—the *endomembrane system*—used to insert membrane proteins, ingest food, and digest cells and macromolecules. Verbal models explaining the origin of endomembranes abound, but explicit quantitative considerations of fitness are lacking. A wealth of quantitative data on vesicle sizes, local protein abundances, protein residence times at functional loci, nutrient transporter rates, membrane protein insertion rates, etc., have been made available in the past couple of decades. Drawing on these data allows for the derivation of two biologically-grounded analytical models of endomembrane evolution that quantify organismal fitness: 1) vesicle-based uptake of nutrient molecules—*pinocytosis*, and 2) vesicle-based insertion of membrane proteins—*proto-endoplasmic reticulum*. Surprisingly, pinocytosis doesn’t provide a net fitness gain under biologically sensible parameter ranges. Explaining why it is primarily used for protein, and not small molecule, uptake in contemporary organisms. The proto-endoplasmic reticulum does provide net fitness gains, making it the more likely candidate for initiating the origin of the endomembrane system. With modifications, the approach developed here can be used to understand the present-day endomembrane system and to further flesh out the cell-level fitness landscape of endomembranes and illuminate the origin of the eukaryotic cell.

## Introduction

Eukaryotic cells are universally endowed with complex and dynamic internal membranes that are topologically separate from the plasma membrane. Examples are the endoplasmic reticulum, Golgi apparatus, endosomes, and transport vesicles^1^. The provenance of these internal membranes, or endomembranes, remains obscure. Many verbal theories of the evolutionary origin of endomembranes have been proposed: endomembranes could have started as plasma membrane attached tubules used for protein excretion^2^, as mitochondrial outer membrane-derived vesicles^3^, or as a phagosome membrane involved in the uptake of prey cells^4,5^. The inside-out model starts with tubules extending outward from what will become the nuclear compartment. These tubules swell into cytoplasmic compartments, and finally the endomembrane system forms where the cytoplasmic compartments meet^6^. Various other works have been published on endomembrane evolution^7–12^.

The general explanatory scheme in these theories is this. A cellular trait of interest is identified in extant species. Combined with the phylogenetic neighbors of these species, an ancestral cellular state is reconstructed. Then a series of intermediates is imagined based on known cell biological principles. The order of all these cellular states then constitutes the evolutionary explanation. This is a helpful approach to understanding the evolution of cell biological traits, but the explanations remain nebulous.

Evolution is a mechanistic process that at base is driven by the population-genetic forces of mutation, recombination, genetic drift, and natural selection^13^. The relative magnitude of these forces determines which phenotypes get access to subsequent generations. It is a numbers game. Quantification is of the essence. Lacking from the verbal theories is an explicit quantitative accounting of the fitness of the cellular states. Preventing us from judging which of these (often incompatible) theories is the real evolutionary sequence of events. Quantitative theories of evolutionary cell biology may help decide between otherwise vague and inconclusive origin stories. Interesting theoretical and quantitative work has been done on later stages of endomembrane evolution^14^ and on the evolution of the nucleus^15^. Presented here is an explicit accounting of the fitness of potential early evolutionary steppingstones toward complex endomembranes.

Two dominant functions of present-day endomembrane systems are (1) the uptake and processing of nutrients, and (2) the production and insertion of membrane proteins. Simplified versions of these processes are clear candidates for intermediates in the evolution of the endomembrane system. For nutrient uptake the simplest system imaginable is a vesicle cycling with transmembrane transport of small-molecule nutrients. For the insertion of membrane proteins, the simplest system is vesicle cycling with membrane protein insertion happening on the vesicles. Both scenarios have endomembranes that are topologically distinct from the plasma membrane.

Key to the calculation of fitness is the observation that eukaryotes exhibit larger cell volumes than prokaryotes^16^. It is likely that the distinguishing features of eukaryotes either enabled this increase in volume or had to evolve in the context of it. As a cell increases its volume and retains its shape, the surface area available per unit volume diminishes. Some processes localized to the plasma membrane don’t need to face the cell surface and can be relegated to internal membranes, freeing up prime real-estate for growth-limiting processes whose access to the external medium is essential.

Presented here are two competing scenarios, with associated mathematical models, of early endomembrane evolution. In both, an intermediate evolutionary stage is considered that consists of a cell in which vesicles emerge from the plasma membrane, spend some time in the cytoplasm, and end their lifetime by fusing back into the plasma membrane. In the pinocytosis scenario, model #1, the vesicles transport nutrient molecules from their lumen into the cytoplasm. In the proto-endoplasmic reticulum scenario, model #2, the vesicles insert membrane proteins into their membrane and deliver these to the plasma membrane upon fusion. For both scenarios the fitness of the derived, vesicle containing, state is calculated with respect to an ancestral state that lacks vesicles. The results have implication for the understanding of the origin of the eukaryotic cell and for understanding the cell biology of extant eukaryotes.

### Defining a fitness function

The pinocytosis and proto-endoplasmic reticulum models utilize the same base function to calculate the net fitness, balancing (1) the gross fitness gain, (2) the energetic cost of introducing a new trait, and (3) the area cost of plasma membrane occupancy by under-construction vesicles.

As a new trait is introduced into the cell, resources are siphoned off from the already existing traits. This causes a reduction in fitness that is approximated as the relative energetic cost of the new trait, *C*_*trait*_, expressed as the ratio between the energetic cost of the new trait divided by the cell budget. Both the trait cost and cell budget are expressed in units of ATP and include opportunity costs^17–19^. The gross fitness gain and the plasma membrane occupancy are combined in a ratio of cell division times of the ancestral (*t*_*d,anc*_) and derived states (*t*_*d,trait*_), explained separately for the two models below. In mathematical form the fitness of the derived state is given by (Supplementary information):

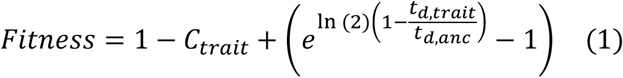

The fitness of the derived state is always evaluated compared to an ancestor of the same cell volume. The ancestral fitness is defined as one.

### Outline of the pinocytosis model

Protein size and rate of conformational change are bounded. For membrane-embedded nutrient transporters, these basic physical facts imply that transport rates over a finite membrane cannot be infinite. Combined with another physical principle, that for expanding cells with invariant shape the cell volume grows faster than the cell surface area, it becomes obvious that large cells face a severe challenge to their growth rate and fitness. If cells grow in volume over evolutionary time, there will come a point at which nutrient demand outstrips transport capacity. Pinocytosis could ameliorate this area-to-volume problem by internalizing some of the external nutrient-containing medium in vesicles and then transporting the nutrients from the vesicle lumens into the cytoplasm, increasing the effective cell surface area for nutrient transport. If the cell growth rate is limited by nutrient transport, this increased nutrient transport capacity increases growth rate and thereby fitness.

The feasibility of pinocytosis as a precursor to the complex endomembrane system is evaluated by comparing two cellular states (Figure 1A). The *ancestral state*—a simple spherical cell that lacks internal membranes and that transports small molecule nutrients over its plasma membrane using nutrient transporters. The *derived state*—a slightly more complex spherical cell that, in addition to having plasma membrane-based nutrient transport, produces vesicles from its plasma membrane that internalize nutrients. With the nutrients being transported from the vesicle lumens into the cytoplasm by vesicle-localized nutrient transporters.

**Figure 1.**
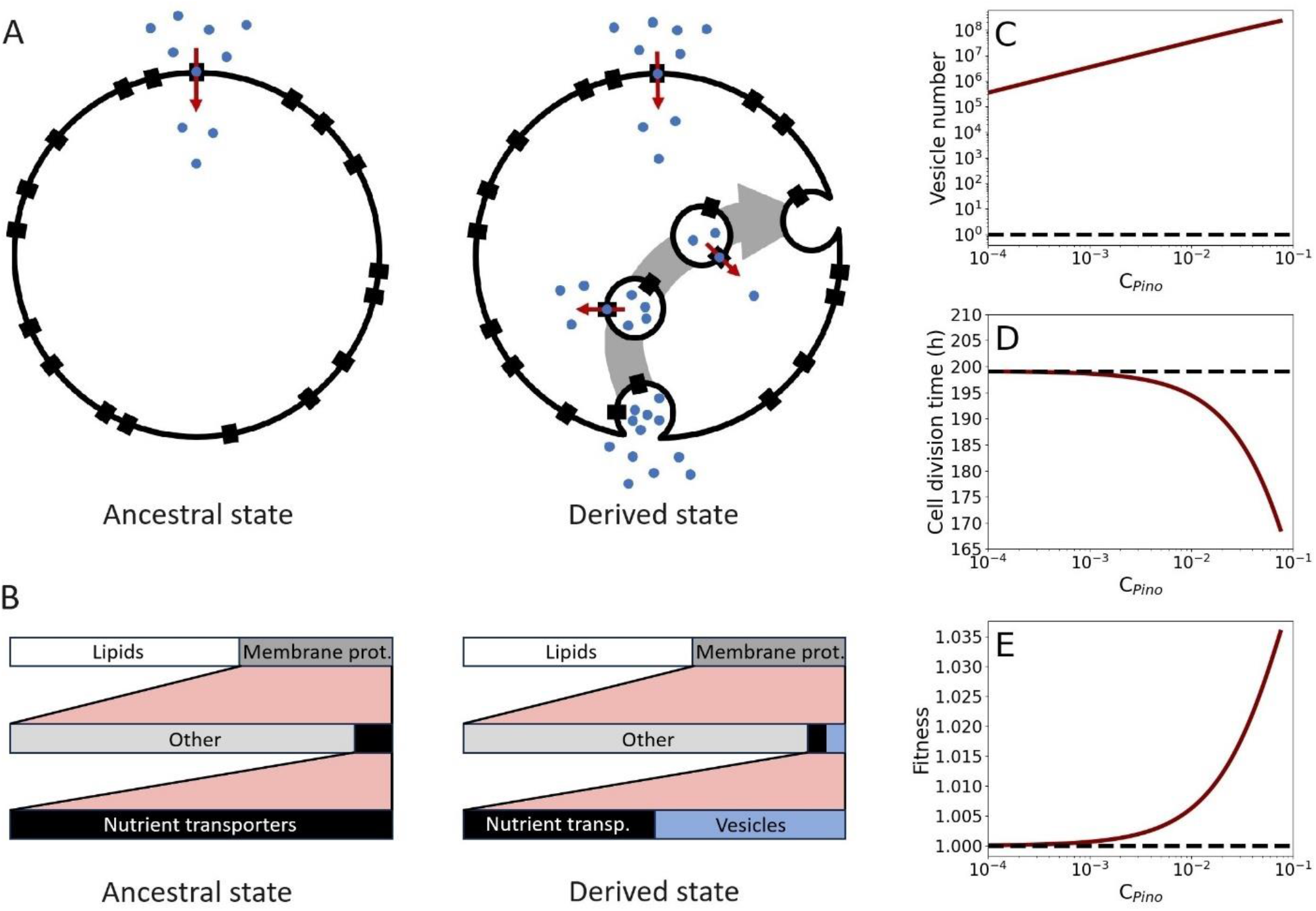
Pinocytosis model and results. (A) Graphical representation of the pinocytosis model, showing the simple ancestral state and the more complex, pinocytic vesicle carrying, derived state. Black rectangles are nutrient transporters, blue dots are nutrient molecules, red arrows show nutrient transport, and the grey arrow shows vesicle progression. (B) Plasma membrane area occupancy in the ancestral and derived state. Length of the bars proportional to fractional occupancy in the plasma membrane. For the derived state the fractional occupancy of under-construction vesicles depends on the investment. (C) Number of vesicles being produced (over the entire cell division period) as a function of the relative investment in pinocytosis. The dashed line shows a vesicle number of one. Cell volume: 10^3^ µm^3^, vesicle radius: 0.025 µm, nutrient concentration: 0.3 M, k_cat_: 1 s^−1^. (D) Cell division time of the derived state as a function of relative investment in pinocytosis. The dashed line shows the ancestral cell division time. Cell volume: 10^3^ µm^3^, vesicle radius: 0.025 µm, nutrient concentration: 0.3 M, k_cat_: 1 s^−1^. (E) Fitness of the derived state as a function of the relative investment in pinocytosis. The dashed line shows the ancestral fitness, normalized to one. Cell volume: 10^3^ µm^3^, vesicle radius: 0.025 µm, nutrient concentration: 0.3 M, k_cat_: 1 s^−1^.

The cell division time of the ancestral state is calculated from the nutrient requirement per unit volume, *N*_*nutV*_, the cell volume at birth *V*_*cell*,0_, and the whole cell nutrient import rate, *v*_*trans,PM*_ (Supplementary information):

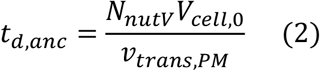

The numerator gives the nutrient requirement for producing a new cell. It is assumed that the cell division time is nutrient limited. The whole cell nutrient import rate is given by:

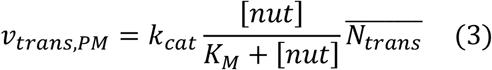

Here, *k*_*cat*_ is the turnover number, [*nut*] is the external nutrient concentration, *K*_*M*_ is the Michaelis constant, and 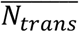 is the number of nutrient transporters averaged over the cell cycle.

For the derived state, the cell division time is calculated similarly but includes the transport rate of nutrients over the combined vesicle membranes, *v*_*trans,pino*_, and discounts the contribution of plasma membrane nutrient transport to provide surface area for vesicle production, *v*_*trans,PM*_^*^ (Supplementary information):

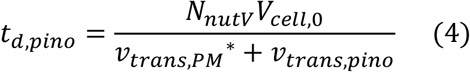

The nutrient transport rate is limited by the area of the plasma membrane and by its composition (Figure 1B). The plasma membrane is assumed to contain 60 % lipids by area, and 40 % protein^20^. Of the protein area, 90 % is assumed to do something other than nutrient transport. In the ancestral state the remaining 10 % of the protein area is used for nutrient transport. In the derived state this 10 % is used both for nutrient transport and for producing vesicles.

The overall nutrient transport rate over the plasma membrane and over the vesicle membranes depends on the number of vesicles present at each moment. The vesicle number is calculated from the investment in pinocytosis, *C*_*pino*_, which is varied to examine its impact on fitness. The vesicle number can only be calculated when the cost per vesicle is known.

### General and pinocytic vesicle costs

Vesicles appear in the models in two classes: the membrane-associated under-construction (non-functional) vesicles and cytoplasmic (functional) vesicles. The cost of under-construction vesicles is the same for both pinocytosis and the proto-endoplasmic reticulum, but the cytoplasmic vesicles have different costs for the two models. For pinocytosis, the cytoplasmic vesicles are 50 nm in diameter and remain the same size over time. The vesicle cost is calculated from the membrane area^21,22^, and accounts for both lipids and membrane proteins. The proto-endoplasmic reticulum cytoplasmic vesicle cost is described in a later section.

In both pinocytosis and the proto-endoplasmic reticulum, vesicles are constructed from the plasma membrane by a dynamic protein assembly, consisting of clathrin, actin, and a slew of other proteins (Table S3). The cost of a single under-construction vesicle is the sum of the average membrane cost and the average cost of the protein assembly that constructs the vesicles (the average being over the construction time of the vesicle). The timing of the stages of vesicle development (Figure S1), the vesicle membrane area over time, and the protein numbers over time are derived from observations in *Schizosaccharomyces pombe*^23^. A detailed breakdown of costs is available in the Supplementary information.

Vesicle fusion is accounted for implicitly by leaving space on the vesicle membrane for fusion proteins such as SNAREs and by the fact that during fusion the vesicle remains functional (the fusion time being subsumed in the cytoplasmic persistence time). The fusion proteins on the plasma membrane are so small in number, compared to the protein assembly required for budding, that they are ignored^24–26^.

### Pinocytosis reduces fitness at low nutrient concentrations

Pinocytosis can only be selected for if nutrient transport over the vesicle membranes can outpace nutrient transport over the plasma membrane. This appears to be the case when a cell is large, the external nutrient concentration is high, and the nutrient transport turnover number, *k*_*cat*_, is low (Figure 1C-E; model parameter values are listed in Table 1). As the investment in pinocytosis is increased, the number of vesicles produced during the cell cycle increases (Figure 1C). This causes the cell division time to decline (Figure 1D) and the fitness to increase (Figure 1E) relative to the ancestor.

**Table 1.** Parameter values used in the models.

| Parameter | Value | Unit | Reference |
| --- | --- | --- | --- |
| Cell cost parameter $C_\alpha$ | $2.7 \times 10^{10}$ | ATP | 18 |
| Nutrient requirement per unit volume | $2 \times 10^9$ | $\mu\text{m}^{-3}$ | 27 |
| $K_M$ for nutrient transport | $10^{-5}$ | M | 53 |
| $k_{\text{cat}}$ for nutrient transport (proto-ER model) | $10^2$ | $\text{s}^{-1}$ | 27–29 |
| Vesicle radius, $r_{\text{ves}}$ | 0.025 | $\mu\text{m}$ | 23 |
| Fraction of the membrane area occupied by protein, $f_p$ | 0.4 | - | 20 |
| Fraction of membrane proteins devoted to other processes, $f_{\text{other}}$ | 0.9 | - | Assumption |
| Area occupancy of membrane protein, $A_{\text{mp}}$ | $10^{-5}$ | $\mu\text{m}^2$ | 27 |
| Volume unit conversion factor, $f_c$ | $10^{-15}$ | $\text{L } \mu\text{m}^{-3}$ | |
| Membrane thickness | 0.005 | $\mu\text{m}$ | 54 |
| Vesicle construction time, $t_{\text{UC}}$ | 140 | s | 23 |
| Vesicle construction time parameter $t_1$ | 110 | s | 23 |
| Vesicle construction time parameter $t_2$ | 10 | s | 23 |
| Vesicle construction time parameter $t_3$ | 20 | s | 23 |
| Vesicle construction length parameter, $l$ | 0.155 | $\mu\text{m}$ | 23 |
| Membrane cost per unit area, $C_{\text{area}}$ | $1.446 \times 10^9$ | $\text{ATP } \mu\text{m}^{-2}$ | 22 |
| External nutrient concentration (proto-ER model) | 0.001 | M | Assumption |
| Membrane protein insertion time, $t_{\text{ins}}$ | 20, 120 | s | See main text. |

The evolutionary importance of pinocytosis can only be judged after examining a wider parameter space. Figure 2 (solid red lines) reveals the fitness at varying cell volume, external nutrient concentration, and turnover number of the nutrient transporter. Only for turnover numbers of 1 s^−1^ and for nutrient concentration of 0.3 M and above, does fitness improve in the derived, pinocytosis exhibiting, state. Larger cell volumes improve the effectiveness of pinocytosis. For the remainder of the tested parameter space the ancestral state has higher fitness. A higher-resolution view of the effect of nutrient concentration and turnover number on fitness leads to the same conclusions (Figure 3A and 3B).

**Figure 2.**
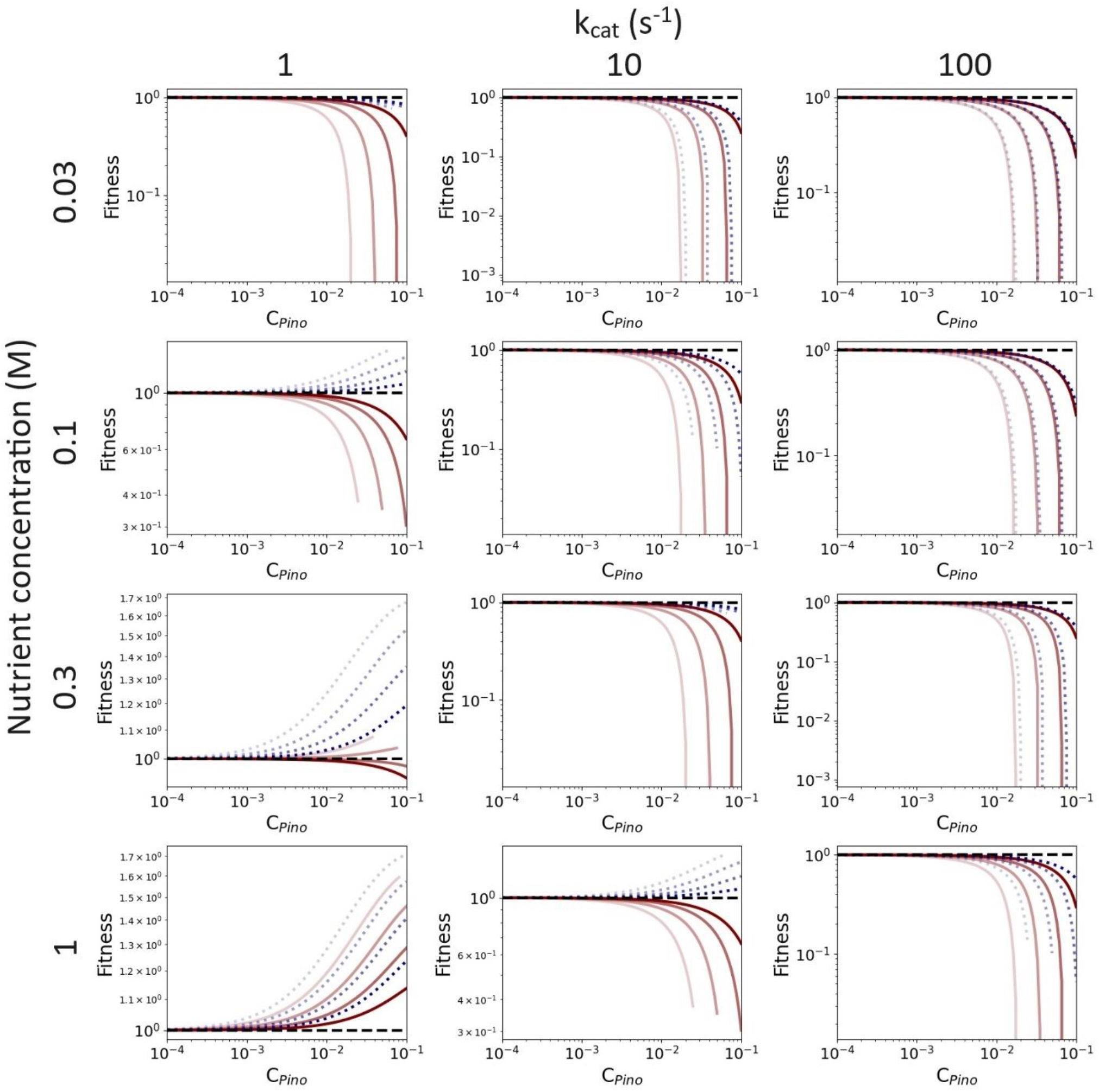
Pinocytosis improves fitness for large vesicles at high nutrient concentration and low transport rate. Fitness of the derived, pinocytosis exhibiting, state as a function of the relative investment of pinocytosis, the volume of the cell, the k_cat_ of the nutrient transporter, the nutrient concentration, and the vesicle size. Black dashed lines are the fitness’ of the ancestral states. Red solid lines: vesicle radius 0.025 µm. Blue dotted lines: vesicle radius 0.1 µm. The cell volume varies from 10, 10^2^, 10^3^, to 10^4^ µm^3^ with the darker lines (for both red and blue) being the smaller volumes. At some level of pinocytosis investment, the under-construction vesicles block the (ancestral) nutrient transporter area on the plasma membrane completely. At this point the model breaks down and the plot is cut off. Cell division times are shown in Figure S2. Vesicle persistence times are listed in Table S1 and S2.

**Figure 3.**
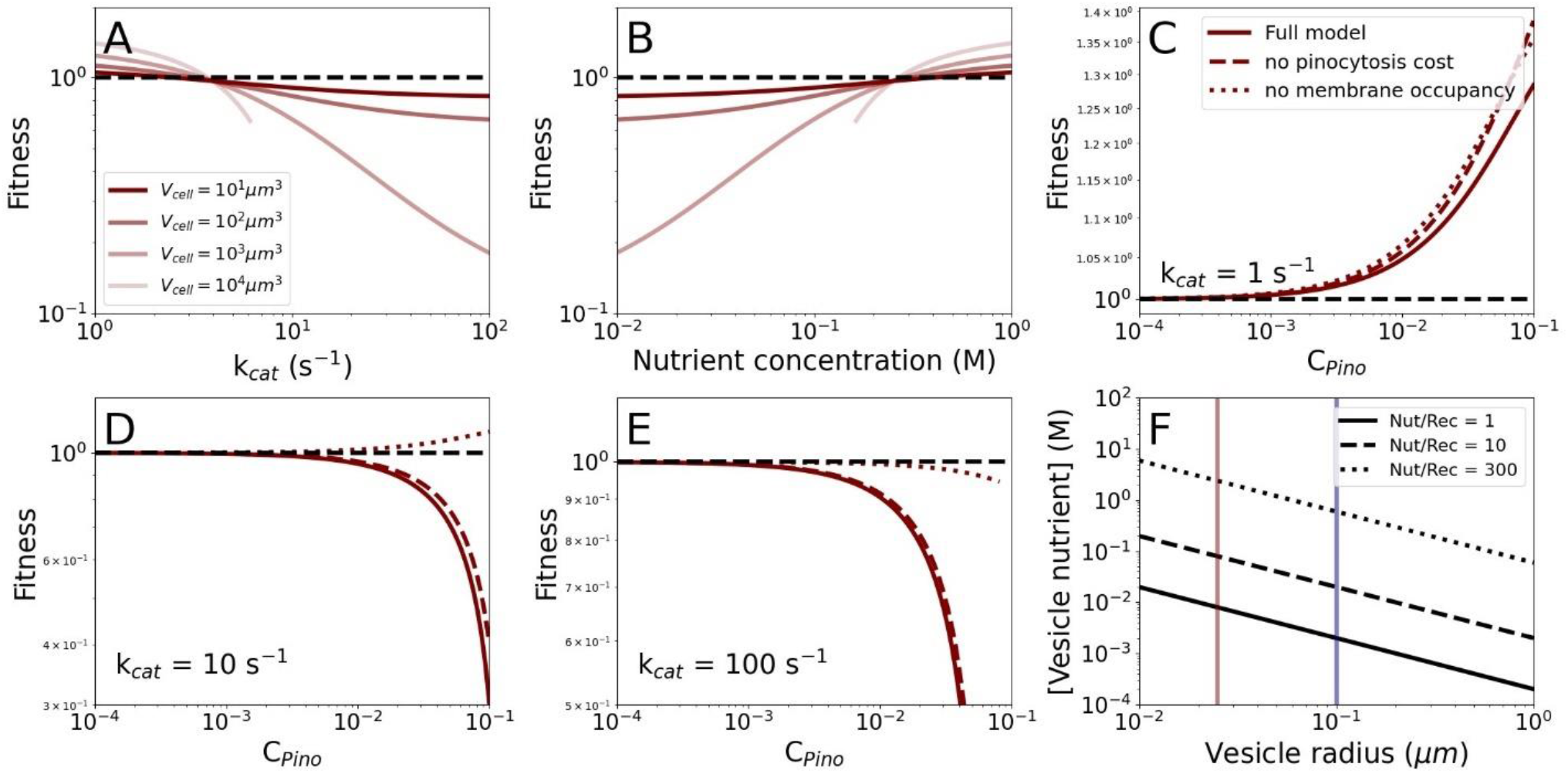
Breakdown of the causes of pinocytosis fitness. (A) Fitness versus the nutrient transport rate parameter k_cat_. Vesicle radius: 0.025 µm, relative pinocytosis cost: 0.03, nutrient concentration: 1 M. For the cell volume of 10^4^ µm^3^, the under-construction vesicles block the nutrient transporter area on the plasma membrane completely at some value of k_cat_. At this point the model breaks down and the plot is cut off. (B) Fitness versus the external nutrient concentration. Vesicle radius: 0.025 µm, relative pinocytosis cost: 0.03, k_cat_: 1 s^−1^. For the cell volume of 10^4^ µm^3^, the under-construction vesicles block the nutrient transporter area on the plasma membrane completely at some nutrient concentration. At this point the model breaks down and the plot is cut off. (C, D, E) Comparison of the fitness calculated from the full model to the fitness calculated in the absence of pinocytosis cost or plasma membrane occupancy by under-construction vesicles. Cell volume: 10^2^ µm^3^, nutrient concentration 1M. (F) The effect of vesicle-localized receptors on internal vesicle nutrient concentration. Vertical lines show the vesicle radii used in the pinocytosis model. Red line: 0.025 µm, Blue line: 0.1 µm.

The turnover number for glucose transport is 100 s^−1^ (reference^27^, see also^28,29^), which means that a cell living on glucose would never evolve pinocytosis to increase nutrient uptake rate. Concentrations of 0.3 M and above are presumably rare in nature. A tightly packed biofilm would have a concentration of nutrients between 1-2 M, but even if our pinocytosing cell were lodged into this biofilm, the nutrients would still need to be freed from the surrounding cells and their macromolecules before they are of use.

A reason for the failure of pinocytosis for increasing nutrient uptake can be gleaned from a breakdown of the constituents of the fitness. There are two detractors to the fitness: (1) the cost of maintaining and producing vesicles and (2) the plasma membrane area occupied by under-construction vesicles. Figures 3C-E show that the largest fitness penalty is the membrane occupation of under-construction vesicles. This means that anything that shortens the residence time, or area occupancy, of under-construction vesicles on the plasma membrane could have a major effect on the possibility of the evolution of pinocytosis.

An increase in vesicle radius could reduce the area occupancy per vesicle-localized nutrient molecule, due to the vesicle volume increasing faster with radius than vesicle cross-sectional area. Vesicle timing and cost have been experimentally determined only for 50 nm diameter vesicles but can be extended to 200 nm diameter vesicles under the following assumptions. The membrane shape remains the same as for the small vesicle throughout vesicle construction, the protein-based construction cost of vesicles increases linearly with vesicle surface area, and the construction time remains the same (Supplementary information). Larger vesicle pinocytosis shows an improvement in fitness over small vesicle pinocytosis (Figure 2), but still depends on low turnover numbers for nutrient transport and high external nutrient concentration.

The requirement for high external nutrient concentration can be ameliorated if the nutrients were concentrated into the vesicles instead of being randomly partitioned between the under-construction vesicle lumen and the external medium. If it is assumed that the entire vesicle surface area available to proteins (40 %) is used to house receptors with a cross-sectional area of 10^−5^ µm^2^, the concentration of nutrients can be calculated for different receptor binding capacities and vesicle radii (Figure 3F). Binding 1 or 10 nutrient molecules to each receptor is insufficient to reach nutrient concentrations high enough for pinocytosis to be effective. It could work if an entire protein’s worth of nutrient molecules, 300 copies, would bind to each receptor. But even this is only good enough when the turnover numbers for nutrient transport are low. Ergo receptors don’t help. Thus, pinocytosis for small molecule nutrient uptake is unlikely to be a precursor to the endomembrane system and is a bad strategy for nutrient uptake in extant organisms.

### Outline of the proto-endoplasmic reticulum model

If a cell volume is increased, the area to volume ratio is reduced. This makes it advantageous to internalize membrane-localized processes that don’t require access to the cell surface, to increase the available area for processes that do need access to the cell surface. The insertion of membrane proteins requires membrane but not direct access to the cell surface. The Sec translocases that facilitate membrane protein insertion, and the associated ribosomes, can thus be housed on internal membranes if a mechanism for membrane protein delivery to the plasma membrane exists^30,31^.

The proto-endoplasmic reticulum model compares the fitness of a simple ancestral cell, lacking internal membranes, to a somewhat more complex cell that houses membrane-bounded vesicles in the cytoplasm (Figure 4A). These vesicles are produced at the plasma membrane and concentrate Sec translocases in their membrane. These Sec translocases insert membrane proteins, causing the vesicles to grow. Lipid insertion into the vesicles is not treated explicitly but is assumed to occur in tandem with membrane protein insertion. After persisting in the cytoplasm for some time these vesicles fuse back into the plasma membrane to deliver their cargo.

**Figure 4.**
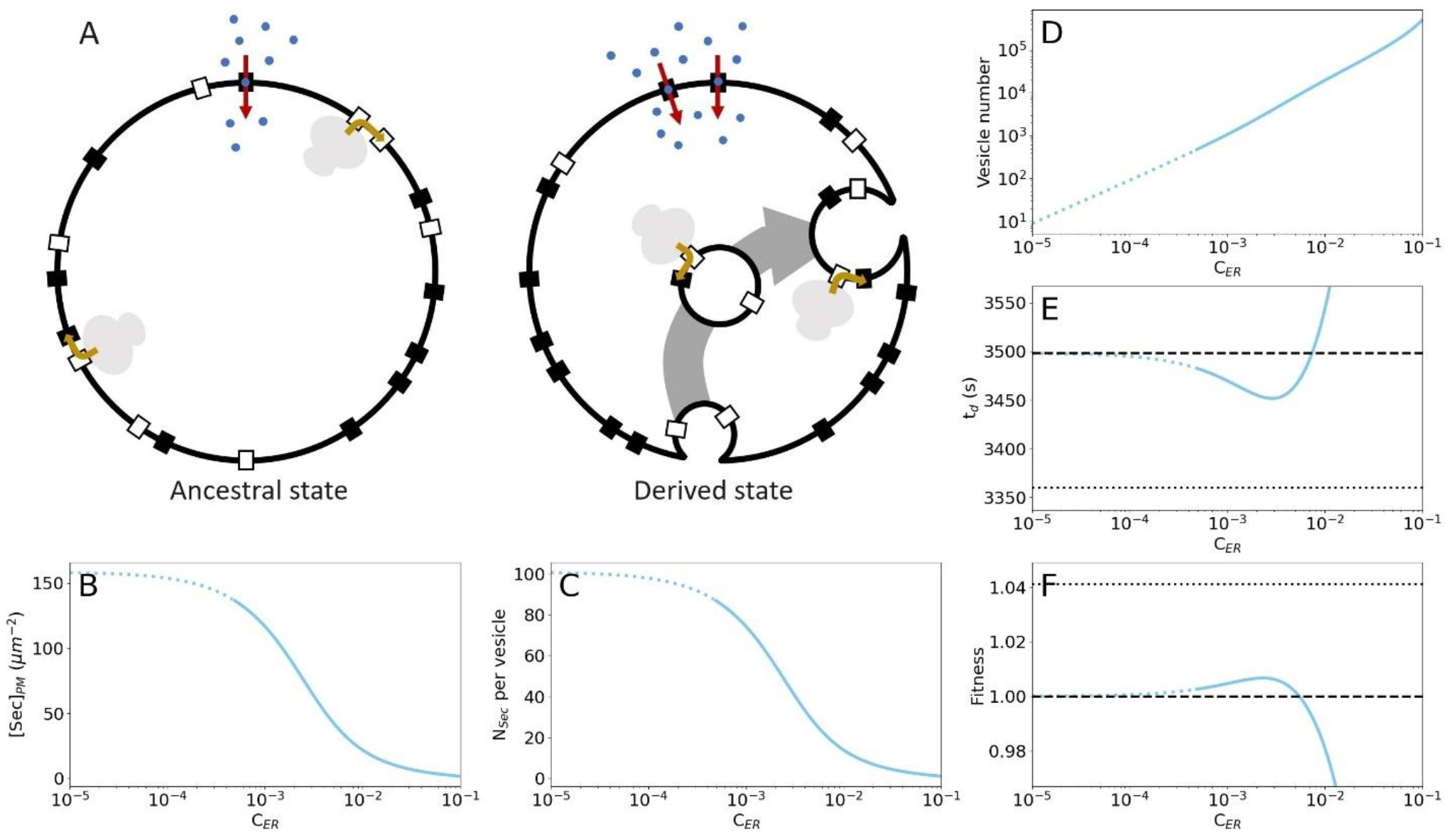
Proto-endoplasmic reticulum model and results. (A) Graphical representation of the proto-endoplasmic reticulum model, showing the simple ancestral state and the more complex, vesicle exhibiting, derived state. Black rectangles are nutrient transporters, white rectangles are Sec translocases, grey bilobed patches are ribosomes, blue dots are nutrient molecules, red arrows show nutrient transport, golden arrows show membrane protein insertion, and the grey arrow shows vesicle progression. (B) Sec translocon concentration in the plasma membrane as a function of the relative cost of the proto-endoplasmic reticulum. (B-F) Membrane protein insertion time (*t*_*ins*_): 20 s, cell volume: 100 µm^3^, vesicle persistence time: 300 s, Sec translocon concentration factor from plasma membrane into vesicles (α): 100. The blue dotted line shows the parameter range over which the model breaks down due to ribosome packing on the vesicle surface. The blue solid line shows where the model works. (C) Number of Sec translocons per vesicle. (D) Total number of vesicles produced throughout the entire cell cycle. (Not the steady state number.) (E) Cell division time of the derived state. Black dashed line: the cell division time of the ancestral state. Black dotted line: the minimal attainable cell division time. (F) Fitness of the derived state. Black dashed line: the fitness of the ancestral state. Black dotted line: the maximal attainable fitness.

This cycling of vesicles permanently internalizes a fraction of the Sec translocases, freeing up area on the plasma membrane for more nutrient transporters. It is this increase in the number of nutrient transporters that via an increased total nutrient import rate and a shorter cell division time (potentially) increases the fitness of the derived state.

The fitness is calculated using Equation 1, the same as for pinocytosis. The difference resides in the calculation of the cell division times. For the proto-endoplasmic reticulum the cell division time of the ancestor is calculated with Equation 2 and 3, but with an 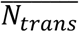 that depends on the number of Sec translocons (Supplementary information). The cell division time of the derived state, *t*_*d,ER*_, is given by:

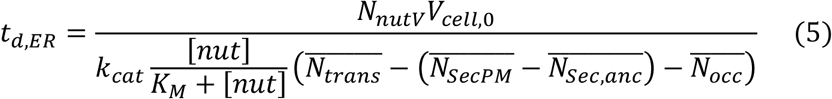

Here 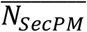 is the number of Sec translocases in the plasma membrane averaged over the cell cycle, 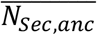 is the average number of Sec translocases in the plasma membrane of the ancestor, and 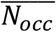 is the average number of nutrient transporters displaced by under-construction vesicles. The combination 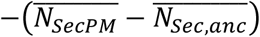 is the gain in the number of nutrient transporters on the plasma membrane. The average numbers of nutrient transporters and Sec translocases^32^ are worked out in the Supplementary information.

### Cytoplasmic vesicle cost in the proto-endoplasmic reticulum

The presence of a proto-endoplasmic reticulum comes at the cost of producing and maintaining the vesicles that constitute it. This cost is calculated in the same way as for pinocytosis, with one exception. In the proto-ER, insertion of membrane proteins (and implicitly lipids) causes the vesicles to grow over time and therefore an average vesicle cost is used. It is assumed that Sec translocase number on the vesicles is constant so that the vesicle surface area grows linearly with time. The details are described in the supplementary information.

### The proto-endoplasmic reticulum improves fitness

As the proto-endoplasmic reticulum is introduced, multiple changes occur in the cell that add up to the change in fitness. Figure 4B shows the concentration of Sec translocon in the plasma membrane being reduced as the investment in the proto-endoplasmic reticulum is increased. This will increase the area on the plasma membrane available for nutrient transporters. The concentration of Sec translocases on each vesicle is determined by applying a fixed concentration factor to the concentration of Sec translocases in the plasma membrane. Thus, the number of Sec translocases per vesicle decreases with investment in the proto-endoplasmic reticulum (Figure 4C). The total number of vesicles produced during the cell cycle increases with investment (Figure 4D), which is what decreases the Sec translocon concentration on the plasma membrane. The changes in the cell shown in Figures 4B-D cause an increase in the number of nutrient transporters which decreases the cell division time (Figure 4E). The cell division time first decreases as the investment in proto-endoplasmic reticulum is increased, but subsequently increases as the excessive production of vesicles competes with nutrient transporters for plasma membrane area. This happens when the area initially devoted to Sec translocases is now completely devoted to production of vesicles. Any increase in the vesicle production after that point will impinge on the area devoted to nutrient transporters. The fitness can be determined by combining the cell division time of the ancestor with that of the derived state and by accounting for the relative construction cost of the proto-endoplasmic reticulum (Equation 1, Figure 4F). The fitness first increases with investment in proto-endoplasmic reticulum and then decreases, by the same mechanism as the cell division time. The one difference being that the fitness gain is somewhat depressed compared to that expected from the cell division times alone because of the cost of producing the vesicles.

As more Sec translocases are loaded onto the vesicles a threshold will be crossed where the ribosomes, which are required for membrane protein insertion, can’t fit onto the vesicle anymore (Supplementary information). Here the model breaks down and the fitness values should be disregarded. The parameter ranges for which this is the case are indicated in the figures.

As with the pinocytosis model the evolutionary relevance can only be judged after scanning the parameter space more thoroughly. In Figure 5 parameter space is scanned over a cell volume of 10^1^-10^4^ µm^3^, a persistence time of 100-1000 s, and a Sec translocon concentration factor of 10-200. This reveals that there is a considerable parameter space over which the proto-endoplasmic reticulum improves fitness. With fitness improvements at larger Sec translocon concentration factors and at longer vesicle persistence times. To simplify the derivation of the model, it was assumed that *t*_*pers*_*/t*_*d*_ ≪ 1 (Supplementary information). In Figure 5 the plots that are faded have a *t*_*pers*_*/t*_*d*_ > 0.1. The consequences of breaking this limit have yet to be examined.

**Figure 5.**
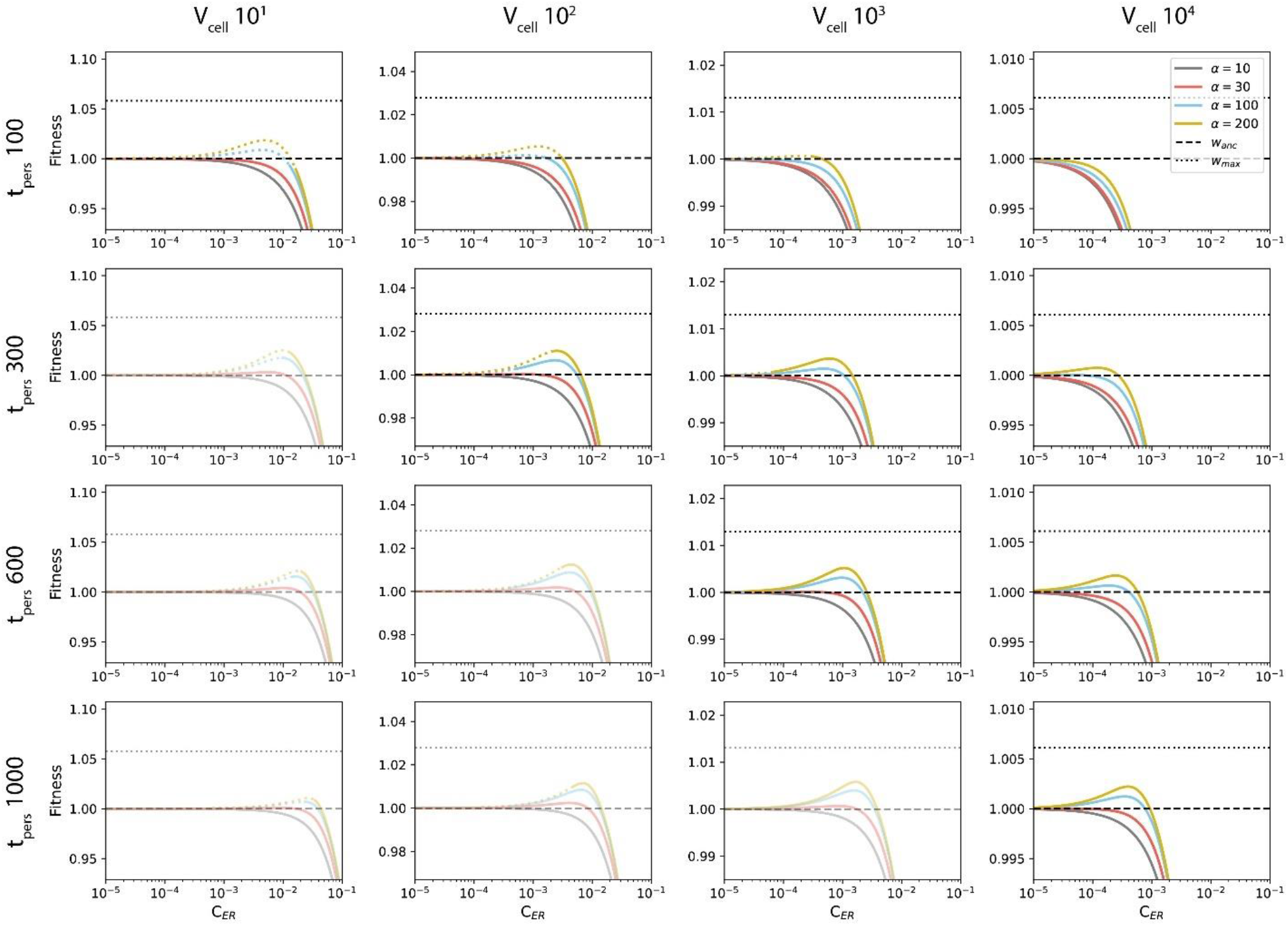
The proto-endoplasmic reticulum improves fitness under reasonable parameter ranges. The fitness of the derived state, containing a proto-endoplasmic reticulum, as a function of the relative cost of the proto-endoplasmic reticulum. The insertion time for individual membrane proteins (*t*_*ins*_) is 20 s. Cell volumes (*V*_*cell*_) in µm^3^, vesicle persistence times (*t*_*pers*_) in seconds. The parameter α shows the concentration factor for Sec translocases from plasma membrane into vesicles. Dotted colored lines show where the model breaks down due to ribosome packing on the vesicle surface. Solid colored lines show where the model works. Dashed black lines: the fitness of the ancestor, dotted black lines: the maximal attainable fitness. Faded plots indicate the parameter combinations for which *t*_*pers*_*/t*_*d*_ > 0.1. To derive the model, it was assumed that *t*_*pers*_*/t*_*d*_ ≪ 1. Cell division times are shown in Figure S3.

Eukaryotes universally carry complex endomembrane systems and have large cells, prokaryotes occasionally have simple endomembrane systems and have small cells. Does the cell volume have an impact on the ease of acquisition of endomembranes? The modeling results are ambivalent (Figure 5). If the *t*_*pers*_*/t*_*d*_ > 0.1 limit is a strict limit, the proto-endoplasmic reticulum works better for cells that are larger than 10 µm^3^, as a wider range of vesicle persistence times show an improvement in fitness for larger volume cells. However, the fitness peaks at a higher value for the smaller cells. This is a direct consequence of the slower growth of the surface area-limited larger cells, as this reduces the required number of Sec translocases as well as the potential gains from internalizing them. The larger fitness gain for smaller cells may increase the rate of fixation of mutants that work toward the proto-endoplasmic reticulum phenotype and thereby improve the chances of evolving the trait for small cells.

The results in Figures 4 and 5 were obtained with a membrane protein insertion time, *t*_*ins*_, of 20 s. This is an estimate derived from the 20 AA/s protein elongation rate in *Escherichia coli*^33^, the average protein length of somewhat over 300 AA^34^, and the knowledge that membrane proteins are inserted co-translationally^30,31^. To account for a potential slowing of protein synthesis during membrane protein insertion, and to account for different protein elongation rates in different species (the lower limit being 3 AA/s^33,35^), the insertion time was extended to 120 s. A longer insertion time leads to higher fitness (Figure 6A and S4).

**Figure 6.**
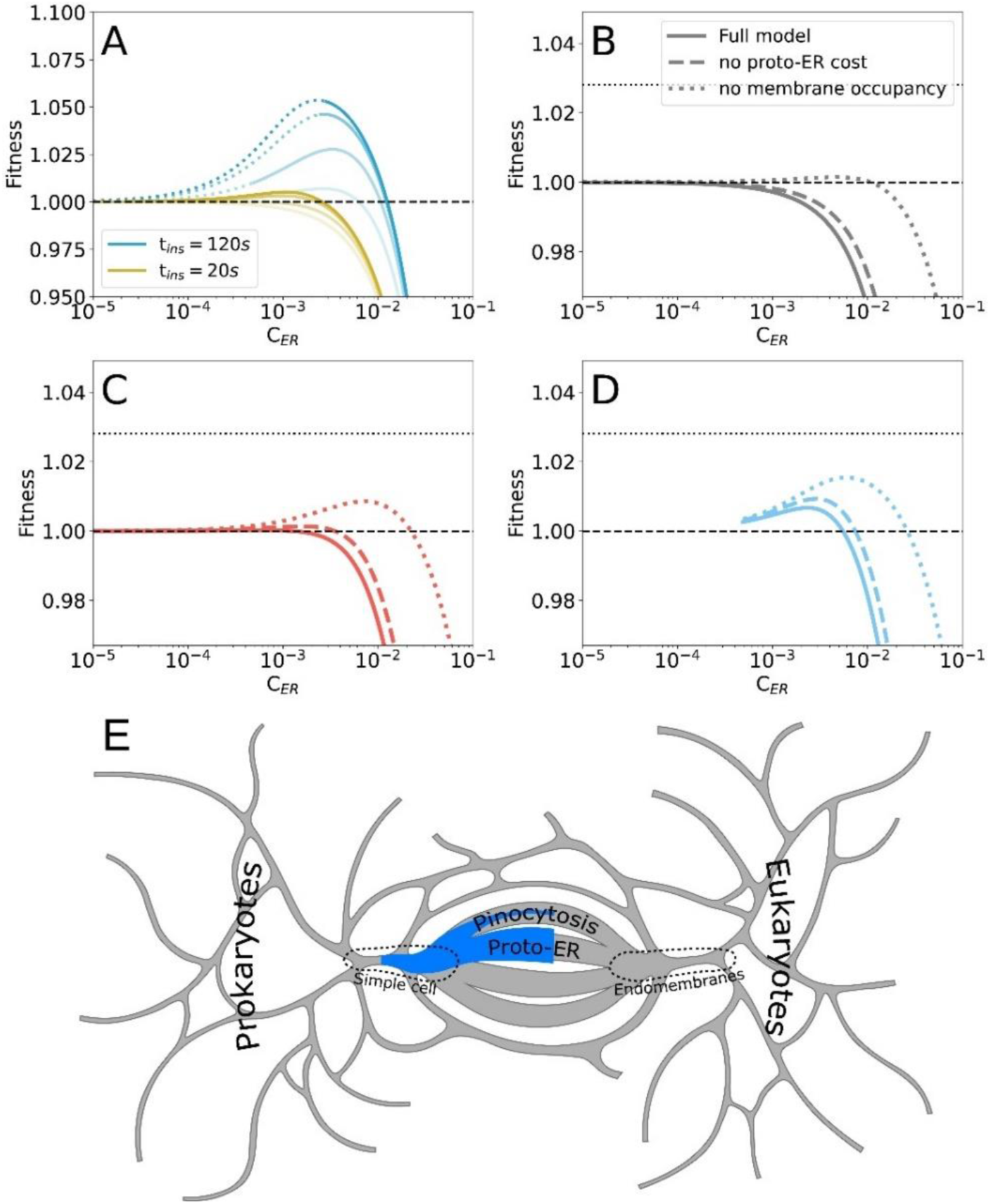
Breakdown of causes of proto-endoplasmic reticulum fitness and cell-level fitness landscape. (A) Fitness of the derived state for two membrane protein insertion times, *t*_*ins*_. The different shades correspond to different concentration factors α, from light to dark 10, 30, 100, and 200. The black dashed line is the ancestral fitness. Cell volume: 10^3^ µm^3^, vesicle persistence time: 600 s. Fitness and cell division times over the full parameter value range are shown in Figure S4 and S5. (B-D) Comparison of the fitness calculated from the full model to the fitness calculated in the absence of pinocytosis cost or plasma membrane occupancy by under-construction vesicles. The black dashed line is the ancestral fitness. The black dotted line is the maximal attainable fitness. The lines in (D) start at the ribosome limit. Cell volume: 10^2^ µm^3^, vesicle persistence time: 300 s, membrane protein insertion time: 20 s. (B) α = 10. (C) α = 30. (D) α = 100. (E) Graphical representation of a highly abstracted cell-level fitness landscape in which each point on the plane is a cellular state associated with a fitness value. White is for cell states that have low fitness and grey is for cell states that have high fitness. The fitness landscape is a hyperdimensional space, here represented in two dimensions for visualization. The branches in the prokaryotic region of the plot correspond to possible prokaryotic cell types, including existing ones. The same is true for the eukaryotic region of the landscape. The grey branching structure shows a potential landscape but at present no meaning should be attached to its actual shape. The blue structure shows the cell-level fitness landscape after applying the models, with the branch thickness indicating the probability of evolution passing through a particular pathway.

Introducing a proto-endoplasmic reticulum comes with two major costs: (1) the energetic costs of producing and maintaining the vesicles, and (2) the cost of occupying plasma membrane area by under-construction vesicles. The contribution of these two effects can be analyzed by simply removing the respective costs from the model and recalculating the fitness. Figures 6B-D show that the plasma membrane area occupation by under-construction vesicles constitutes the largest penalty, as it was for the pinocytosis model.

### An intuitive explanation for the pinocytosis and proto-ER models

A patch of plasma membrane area can only be used by one process at a time. For pinocytosis to work, the benefit of producing a vesicle must outstrip the benefit of using that same patch of plasma membrane area for nutrient transport directly. The maximal number of nutrient molecules that can be imported into the cytoplasm by a single vesicle is equal to the number of nutrient molecules present in the vesicle just after it is formed. And it is this number that needs to exceed the number of nutrients being transported directly if this patch of plasma membrane area was occupied by nutrient transporters. This explains why larger vesicles and higher external nutrient concentrations work better: they provide more nutrient molecules per vesicle per unit plasma membrane area. This also explains why a slower nutrient transporter works in favor of pinocytosis: it lowers the threshold number of nutrient molecules that are required to be present in the pinocytic vesicles at the start.

The proto-endoplasmic reticulum faces a similar trade-off. The benefit of producing a vesicle needs to outstrip the benefit of using that same patch of membrane area for membrane protein insertion. However, the proto-endoplasmic reticulum has two crucial advantages over pinocytosis. First, the number of membrane protein insertions that a single vesicle can accomplish increases with the time the vesicle spends in the cytoplasm. This is different from the pinocytosis model where no further gains can be made after the vesicle is emptied. The second advantage of the proto-endoplasmic reticulum model is that the extra nutrient transporters are added to the plasma membrane where the pool of nutrients is effectively infinite. In the pinocytosis model, in contrast, the extra nutrient transporters are added on the vesicles where the pool of nutrient molecules is small.

The pinocytosis model does have one advantage over the proto-endoplasmic reticulum model: the potential spoils are much larger. The maximal fitness gains for the proto-endoplasmic reticulum are limited by the relatively small Sec translocon requirement in the ancestor. The pinocytosis model doesn’t suffer from this limitation, as is evident from Figure 2. Overall, however, it appears that the proto-endoplasmic reticulum model succeeds under sensible biological parameters, whereas the pinocytosis model works only under excessively high nutrient concentrations. Therefore, a larger weight can be assigned to an evolutionary path toward complex endomembranes that passes through a proto-endoplasmic reticulum.

### Eukaryogenesis and a cell-level fitness landscape

Cell biological transitions are the result of thousands of mutations and fixation events. To guide our understanding of this evolutionary process, we require a coarse-grained understanding of the fitness contributions associated with intermediate states along evolutionary paths. The verbal theories alluded to in the introduction^2–12^ provide such a coarse-grained understanding, and when combined these constitute some kind of fitness landscape where each position is a cellular state (Figure 6E). Our approach is complementary to the verbal theories by applying physical principles, and empirically determined parameter values, to explicitly assign a fitness to these cellular states. These act as weights (or probabilities) on the evolutionary paths (Figure 6E, blue network). Pinocytosis of small molecule nutrients is unlikely to work in natural environments and receives a smaller weight, whereas the proto-endoplasmic reticulum does work under natural conditions and therefore receives a larger weight. Ultimately, this approach can be extended to the evolution of endomembrane-based protein excretion, membrane homeostasis, protein glycosylation, and even the cytoskeleton and the nucleus, to help understand eukaryogenesis and the evolution of cell types more generally.

### Why is pinocytosis present in extant organisms?

Pinocytosis is the vesicle-based uptake of fluid and solutes. According to the pinocytosis model described above it is unlikely that the solutes are small molecule nutrients. This is borne out by a survey of the pinocytosis literature. *Amoeba proteus* performs pinocytosis via channels that pinch of 1-2 µm diameter vesicles at the tip^36^, much larger than what we studied, and though uptake of sucrose by pinocytosis has been observed in this species^37^, it is not shown that this is the nutrient source. Other studies in amoebae describe pinocytosis for protein uptake^38^. Uptake of vitamin B12 by pinocytosis in rat epithelial cells occurs only when the vitamin is bound to the protein gastric intrinsic factor^39^. A description of pinocytosis in *Acanthamoeba* examines only protein uptake^40^. The peripheral vesicles in *Giardia* play a role in pinocytic uptake of proteins^41^. The cytostome-cytopharynx complex in *Trypanosoma* is used for vesicle-based uptake of solutes, but again only protein uptake is described^42^. Micropores in gregarine apicomplexans may be used for pinocytosis but it is not specified what the solutes are^43^. The parabasalid *Tritrichomonas* exhibits pinocytic vesicles, but only protein uptake is examined^44^. Amoebae and animals perform macropinocytosis, the uptake of fluid and solutes in large vesicles. It is suggested to have evolved for food uptake in the form of proteins or other macromolecules^45^. In animal cells, pinocytosis appears to be used for uptake of macromolecules and lipid-protein particles^46^. In contrast to all these species that perform pinocytosis, fungi digest their food externally and take up small molecules, but uptake is facilitated by plasma membrane localized transporters^47^. Concluding, it appears that pinocytosis is used in many species across eukaryote diversity, but that the target solutes are proteins, not small molecules. And in the major group of eukaryotes known to import small molecule nutrients, the fungi, this is performed by plasma membrane localized transporters, not pinocytosis.

### Extension to other endomembrane-based processes

Pinocytosis and a proto-endoplasmic reticulum are distinct biological phenomena but, as demonstrated in this work, can be understood via similar models. This has implications for cell biology more broadly. A plethora of endomembrane-associated traits in extant organisms depend on the transfer of vesicles. And an understanding of these phenomena requires a comparison with alternative cellular architectures that lack such vesicle trafficking, as is done here for pinocytosis and the proto-endoplasmic reticulum. Lipid production^48^ and protein glycosylation^49^ are like membrane protein insertion in that the enzymes involved occupy space on a membrane. If we know the rates and sizes of the involved enzymes, the internalization of lipid production and glycosylation can be described by the proto-endoplasmic reticulum model. Protein secretion could also be described like the proto-endoplasmic reticulum, if an additional component is included that accounts for the fitness benefit of excreted protein. This could be compared to a model in which proteins are excreted via membrane attached tubules, as described verbally before^2^. Membrane proteins in the plasma membrane may get oxidized or misfold and therefore require quality control, the removal of bad proteins and delivery of new proteins is facilitated by vesicle exchange^50^. The same holds true for the swapping out of different types of membrane proteins in response to a change in the environment^51^. Both recycling and membrane reconfiguration can alternatively be accomplished by having Sec translocases and membrane bound proteases (FtsH in *Escherichia coli*^52^) on the plasma membrane, as is the case in bacteria. Transport of proteins and lipids between endomembrane compartments, from endoplasmic reticulum to Golgi, between Golgi stacks, from Golgi to endosomes, etc.^1^, follows similar principles as the pinocytosis and proto-endoplasmic reticulum model but presumably doesn’t suffer the same membrane area constraint because additional membrane can easily be added. Combining these considerations, we may progress on the deeper question of why ubiquitous vesicle transport exists in the first place, and why evolution decided against putting a dedicated Sec translocase on every internal compartment.

### Membrane tubules and cell elongation as alternatives to vesicles

Both cell elongation and plasma membrane-attached internal membrane tubules^2^, by increasing the area-to-volume ratio of the cell, could be used to achieve the same goal as pinocytosis and the proto-endoplasmic reticulum. The reasons for choosing for vesicles over tubules are twofold: (1) detailed experimental studies were available that quantify the requirements of vesicle construction, and (2) the vesicles are a separate compartment and therefore closer to the eukaryote endomembrane system. When the cost of tubule construction and maintenance are better understood, a proper comparison between vesicles and tubules can be performed. Vesicles do have the advantage that the proteins that determine their structure need to exist only during the construction phase, whereas maintaining tubule shape requires a permanent presence of the requisite proteins. Cell elongation has fitness consequences and can’t be pursued indefinitely. Our models can still be applied to the somewhat elongated cells at the point where the cells can’t be made longer without compromising fitness. This would yield similar results but shifted with respect to cell volume.

## Conclusion

Two competing models for the early evolution of endomembranes were derived in which fitness is calculated explicitly from cell biological traits. The pinocytosis model fails to improve fitness under biological plausible nutrient concentrations. The proto-endoplasmic reticulum model doesn’t suffer from this problem and is the more likely transitional state from simple cells, that lack internal membranes, to modern eukaryotic cells that sport a manifold endomembrane system. Surprisingly, the uptake of small molecules by pinocytosis isn’t favored by natural selection, but this finding is consistent with, and explains, the distribution and nature of pinocytosis across the tree of life. Extensions of the approach presented here will improve the understanding of eukaryotic origins and the cell biology of present-day organisms.

## Acknowledgements

We thank Noah Spencer, John McCutcheon, and Jeremy Wideman for commenting on a draft of the paper. This work was funded by the Moore–Simons Project on the Origin of the Eukaryotic Cell, Simons Foundation 735927 (https://doi.org/10.46714/735927), the National Institutes of Health, R35-GM122566-01, and the National Science Foundation, DBI-2119963.

