## Supplementary information for "Quantifying the evolutionary paths to endomembranes"

### Defining a fitness function

The fitness of the derived state is determined from the cell division times of the ancestral and derived states and by accounting for the cost of performing pinocytosis. Using a discrete generations model the fitness of the derived state is:

$$Fitness = 1 - s_{cost} + s_{gain} \qquad (S1)$$

Here $s_{cost}$ and $s_{gain}$ are selection coefficients. Building on earlier work[5], $s_{cost}$ is equated to the relative energetic cost of the introduced trait, $C_{trait}$ (compared to the cell budget). The gain depends on the cell division times of the ancestral and derived states. Cell division times are properties of a continuous growth model and are therefore converted to align with the fitness equation above. Comparing a discrete growth model for the ancestor to a continuous one,

$$N_{anc} = N_{anc,0} a^{\frac{t}{\Delta t}} \qquad (S2)$$

$$N_{anc} = N_{anc,0} e^{\frac{\ln(2)}{t_{d,anc}} t} \qquad (S3)$$

we obtain:

$$a^{\frac{1}{\Delta t}} = e^{\frac{\ln(2)}{t_{d,anc}}} \qquad (S4)$$

Here $N_{anc}$ is the number of ancestor cells after time $t$, $N_{anc,0}$ is the number of ancestor cells at time zero, $a$ is the growth rate parameter, $\Delta t$ is the generation time, and $t_{d,anc}$ is the ancestral cell division time. Equation S4 can be rewritten to:

$$a = e^{\frac{\ln(2)}{t_{d,anc}} \Delta t} \qquad (S5)$$

For the derived state the growth models are:

$$N_{der} = N_{der,0} a(1+s)^{\frac{t}{\Delta t}} \qquad (S6)$$

$$N_{der} = N_{der,0} e^{\frac{\ln(2)}{t_{d,der}} t} \qquad (S7)$$

This yields:

$$a(1+s)^{\frac{1}{\Delta t}} = e^{\frac{\ln(2)}{t_{d,der}}} \qquad (S8)$$

Here $s$ is a selection coefficient and $t_{d,der}$ is the cell division time of the derived state. Equation S8 can be rewritten to:

$$s = \frac{e^{\frac{\ln(2)}{t_{d,der}} \Delta t}}{a} - 1 \qquad (S9)$$

Substituting Equation S5 yields:

$$s = \frac{e^{\frac{\ln(2)}{t_{d,der}}\Delta t}}{e^{\frac{\ln(2)}{t_{d,anc}}\Delta t}} - 1 = e^{\ln(2)\left(\frac{\Delta t}{t_{d,der}} - \frac{\Delta t}{t_{d,anc}}\right)} - 1 \qquad (S10)$$

The generation time, $\Delta t$, must be chosen such that it matches with the generation time associated with $s_{cost}$. This is the cell division time of the derived state, $t_{d,der}$, as the costs are not present in the ancestor. The fitness function then becomes:

$$Fitness = 1 - C_{trait} + \left(e^{\ln(2)\left(1 - \frac{t_{d,der}}{t_{d,anc}}\right)} - 1\right) \qquad (S11)$$

### Pinocytosis model

The pinocytosis model compares two cellular states. A simple ancestor that transports nutrients over its plasma membrane by transmembrane transporters, and a more complex derived state that in addition to plasma membrane-based nutrient transport produces vesicles from its plasma membrane. The vesicle lumens contain nutrients that are transported into the cytoplasm by vesicle membrane-localized nutrient transporters. After persisting for some time in the cytoplasm, the vesicles fuse back into the plasma membrane. The purpose of pinocytosis is to increase the effective cell surface area so that an otherwise surface-area limited cell can increase its growth rate by increasing nutrient transport rate.

#### Pinocytosis—Cell division time of the ancestral state

The cell division time of the ancestor is set by the rate of nutrient uptake relative to the nutrient requirement for producing a new cell.

$$t_{d,anc} = \frac{N_{nutV} V_{cell,0}}{v_{trans,PM}} \qquad (S12)$$

With $N_{nutV}$ being the nutrient requirement per unit volume, $V_{cell,0}$ being the cell volume at the start of the cell cycle, and $v_{trans,PM}$ being the nutrient transport rate over the entire plasma membrane. The transport rate is determined by Michaelis-Menten kinetics, and by the total number of nutrient transporters present on the plasma membrane.

$$t_{d,anc} = \frac{N_{nutV} V_{cell,0}}{k_{cat} \frac{[nut]}{K_M + [nut]} \overline{N_{trans}}} \qquad (S13)$$

With $k_{cat}$ and $K_M$ being the standard enzyme parameters, $[nut]$ being the external nutrient concentration, and $\overline{N_{trans}}$ being the average number of transporters present on the plasma membrane over the cell cycle. The transport rate of nutrient molecules from bulk medium to the cell surface is assumed to be significantly faster than the transport rate over the membrane, so that diffusion rates don't need to be accounted for explicitly. The number of nutrient transporters is related to the cell volume by:

$$\overline{N_{trans}} = \frac{N_{trans,0}}{ln(2)} = \frac{A_{pm,0} f_p (1 - f_{other})}{A_{mp} ln(2)} = \frac{4\pi \left(\frac{3V_{cell,0}}{4\pi}\right)^{\frac{2}{3}} f_p (1 - f_{other})}{A_{mp} ln(2)} \qquad (S14)$$

Here, $N_{trans,0}$ is the number of nutrient transporters present at the start of the cell cycle, $A_{pm,0}$ is the plasma membrane area at the start of the cell cycle, $f_p$ is the fraction of the membrane area that is protein, $f_{other}$ is the fraction of other membrane proteins (in the pinocytosis model this means every membrane protein that is not a nutrient transporter), $A_{mp}$ is the surface area on the membrane occupied by a single membrane protein (all membrane proteins are assumed to have the same area occupancy), and $V_{cell,0}$ is the cell volume at the start of cell division. The cell is assumed to be spherical at the start of the cell cycle so that $A_{pm,0} = 4\pi\left(\frac{3V_{cell,0}}{4\pi}\right)^{\frac{2}{3}}$. As the cell cycle progresses the cell elongates to keep the area to volume ratio constant during the cell cycle, which obviates the need for a burst of membrane production at cell division (due to area/volume scaling) and simplifies the model.

The relation between $\overline{N_{trans}}$ and $N_{trans,0}$ is explained in the "Correcting for cell growth" section.

**Pinocytosis—Cell division time of the derived state**

The derived state has nutrient transporters on the plasma membrane just like the ancestral state, but in addition has vesicles budding from the plasma membrane. These vesicles persist for some time in the cytoplasm and then fuse back into the plasma membrane. As the vesicles form, they internalize some of the external fluid containing the nutrients. The nutrients are transported from the vesicle lumen into the cytoplasm by vesicle-localized nutrient transporters. The formation of vesicles blocks some of the plasma membrane area from having nutrient transporters, so the rate of nutrient transport over the plasma membrane is reduced in the derived state compared to the ancestral state. The cell division time of the derived state is given by:

$$t_{d,pino} = \frac{N_{nutV}V_{cell,0}}{v_{trans,PM}{}^{*} + v_{trans,pino}} \qquad (S15)$$

Here, $v_{trans,PM}{}^{*}$ is the adjusted nutrient transport rate over the entire plasma membrane and $v_{trans,pino}$ is the nutrient transport rate over all the pinocytic vesicles in the cell.

Each pinocytic vesicle starts with a nutrient concentration equal to the external nutrient concentration. The nutrients are transported from the vesicle lumen into the cytoplasm by vesicle membrane-localized nutrient transporters. The average transport rate for each vesicle is set by the total number of nutrient molecules present in the vesicle and the time the vesicle spends in the cytoplasm (the persistence time, $t_{pers}$):

$$v_{trans,pino} = \frac{N_A f_c V_{lumen}[nut]N_{ves,0}}{t_{pers}ln(2)} \qquad (S16)$$

Here, $N_A$ is Avogadro's number, $f_c$ is a conversion factor for the volume units (vesicle volume is in μm$^3$ and nutrient concentration in mol L$^{-1}$), $V_{lumen}$ is the vesicle lumen volume obtained from $V_{lumen} = \frac{4\pi}{3}(r_{ves} - h_{mem})^3$ with $h_{mem}$ being the thickness of the membrane, $[nut]$ is the external nutrient concentration, $N_{ves,0}$ is the number of vesicles in the cell at the start of the cell cycle ($\frac{N_{ves,0}}{ln(2)}$ is the average number of vesicles present in the cell over the cell cycle), and $t_{pers}$ is the persistence time of the vesicle. The persistence time is calculated from the nutrient and transporter concentrations in the vesicle:

$$t_{pers} = \frac{[nut]}{k_{cat}[trans]} = \frac{[nut]}{k_{cat}\frac{4\pi\left(r_{ves} - \frac{h_{mem}}{2}\right)^2 f_p(1 - f_{other})}{A_{mp}V_{lumen}N_A f_c}} \qquad (S17)$$

Here, $[nut]$ is the external nutrient concentration, $[trans]$ is the nutrient transporter concentration in the entire volume of the vesicle, $h_{mem}$ is the membrane thickness, and $V_{lumen}$ is the vesicle lumen volume. To calculate $t_{pers}$ it is assumed that $[nut] \gg K_M$, so that the transport rate reduces to $k_{cat}$.

The number of vesicles at the start of the cell cycle, $N_{ves,0}$, is calculated from the investment in pinocytosis and is calculated in the "Obtaining vesicle number from pinocytosis investment" section.

The adjusted nutrient transport rate over the plasma membrane is given by:

$$v_{trans,PM}^{*} = k_{cat}\frac{[nut]}{K_M + [nut]}\left(\overline{N_{trans}} - \overline{N_{occ}}\right) \qquad (S18)$$

With $\overline{N_{trans}}$ being the average number of transporters on the plasma membrane of the ancestral state, and $\overline{N_{occ}}$ being the average number of transporters that are absent because the area they would have occupied is now used to produce vesicles. The number of transporters that is displaced by the vesicles is explained in the "Pinocytosis vesicle cost" section, and is given by:

$$\overline{N_{occ}} = \frac{\pi r_{ves}^2\left(\frac{1}{2}\frac{t_1}{t_{pers}} + \frac{t_2}{t_{pers}}\right) f_p N_{ves,0}}{A_{mp} ln(2)} \qquad (S19)$$

Here, $t_1$, $t_2$, and $t_{UC}$ are time periods in the construction of vesicles, explained in the "Pinocytosis vesicle cost" section. The number of under-construction vesicles is given by $\frac{t_{UC}}{t_{pers}}N_{ves,0}$, where it is assumed that the vesicle construction time, $t_{UC}$, and the vesicle persistence time, $t_{pers}$, are short enough relative to the cell division time so that the cell growth effect on (under-construction) vesicle numbers can be ignored.

**Pinocytosis vesicle cost**

Vesicles appear in the pinocytosis and proto-endoplasmic reticulum models in two classes: the membrane-bound under-construction (non-functional) vesicles and cytoplasmic (functional) vesicles. Because vesicles are produced and removed at fixed rates (relative to the cell surface area), the investment in vesicles is constant relative to the cell budget. The magnitude of this investment can then be obtained by examining the vesicle abundance at a single time point in the cell cycle. Throughout the calculations this timepoint is taken to be the start of the cell cycle.

The cost of individual cytoplasmic vesicles is just the membrane cost (in units of ATP, and including opportunity costs)[1,2], and accounts for both lipids and proteins. With proteins constituting 40% of the membrane area[3]. The cytoplasmic vesicles have a diameter of 0.05 µm, and their cost is given by:

$$C_{ves,0} = 4\pi\left(r_{ves} - \frac{h_{mem}}{2}\right)^2 C_{area} N_{ves,0} \qquad (S20)$$

Here, $C_{ves,0}$ is the total cytoplasmic vesicle cost in the cell at the start of the cell cycle (in ATP), $r_{ves}$ is the vesicle radius, $h_{mem}$ is the membrane thickness, $C_{area}$ is the cost (in ATP) per μm$^2$ of membrane, and $N_{ves,0}$ is the number of cytoplasmic vesicles present in the entire cell at the start of the cell cycle.

The cost of the under-construction vesicles is based on detailed quantitative microscopy of *Schizosaccharomyces pombe*[4] and includes the cost of the membrane and the cost of the proteins that assemble on the plasma membrane and construct the vesicle. The vesicle forms in three morphologically distinct stages: 1) the transition from a flat surface to a membrane-attached hemisphere, 2) the transition from a hemisphere to a pillar (a cylinder capped with a hemisphere), and 3) the persistence of a non-functional spherical vesicle at the membrane (Figure S1). The average area of the under-construction vesicle is given by:

$$\overline{A_{UC}}^{*} = \frac{2\pi\left(r_{ves} - \frac{h_{mem}}{2}\right)^2}{2}\frac{t_1}{t_{UC}} + \left(2\pi\left(r_{ves} - \frac{h_{mem}}{2}\right)^2 + \pi\left(r_{ves} - \frac{h_{mem}}{2}\right)l\right)\frac{t_2}{t_{UC}} + 4\pi\left(r_{ves} - \frac{h_{mem}}{2}\right)^2\frac{t_3}{t_{UC}} \qquad (S21)$$

Here, $r_{ves}$ is the radius of the vesicle, $h_{mem}$ is the membrane thickness, $t_1$, $t_2$, or $t_3$ is the time the vesicle spends in stage 1, 2, or 3, $t_{UC}$ is the overall construction time, and $l$ is the length of the cylinder in stage 2. It is assumed that addition of membrane to the hemisphere of stage 1 and the cylinder of stage 2 proceeds linearly with time, allowing the calculation of the average membrane area by dividing the area of the complete hemisphere (in stage 1) and cylinder (in stage 2) by two. Because the vesicle membrane is formed out of, and connected to, the plasma membrane, there is a patch of plasma membrane missing which reduces the cost of the under-construction vesicle. Accounting for this yields:

$$\overline{A_{UC}} = \frac{\pi\left(r_{ves} - \frac{h_{mem}}{2}\right)^2}{2}\frac{t_1}{t_{UC}} + \left(\pi\left(r_{ves} - \frac{h_{mem}}{2}\right)^2 + \pi\left(r_{ves} - \frac{h_{mem}}{2}\right)l\right)\frac{t_2}{t_{UC}} + 4\pi\left(r_{ves} - \frac{h_{mem}}{2}\right)^2\frac{t_3}{t_{UC}} \qquad (S22)$$

This is the area used to calculate the membrane cost of under-construction vesicles in the models. The cost is obtained using:

$$C_{UC,0} = \overline{A_{UC}}C_{area}\frac{t_{UC}}{t_{pers}}N_{ves,0} \qquad (S23)$$

Where it is assumed that the vesicle construction time, $t_{UC}$, and the vesicle persistence time, $t_{pers}$, are short enough relative to the cell division time so that the cell growth effect on (under-construction) vesicle numbers can be ignored. The number of under-construction vesicles is given by $\frac{t_{UC}}{t_{pers}}N_{ves,0}$.

The vesicles are extracted from the plasma membrane by the combined action of 16 proteins (Table S3), including clathrin and actin, that change in abundance over the construction period, $t_{UC}$ [4]. The cost of the vesicle producing proteins is the product of the number of under-construction vesicles, the average number of proteins of each type over the construction period, $t_{UC}$, and the cost per individual protein (in ATP). This yields:

$$C_{prot,0,i} = \frac{t_{UC}}{t_{pers}} N_{ves,0} \times \frac{N_{pp,i}}{2} \frac{t_{prot,i}}{t_{UC}} \times N_{AA,i} C_{AA} \qquad (S24)$$

Here, $C_{prot,0,i}$ is the combined cost of all the monomers of protein $i$ present on all vesicles, $N_{pp,i}$ is the number of monomers of protein $i$ present at the peak abundance, $t_{prot,i}$ is the residence time of protein $i$ at the under-construction vesicle, $N_{AA,i}$ is the number of amino acids in protein $i$, $C_{AA}$ is the average cost of a single amino acid, and $\times$ is an ordinary product (not a cross product). This can be simplified and summed for all 16 proteins:

$$C_{prot,0} = \frac{N_{ves,0} C_{AA}}{2 t_{pers}} \sum_{i}^{All} N_{pp,i}\, t_{prot,i} N_{AA,i} \qquad (S25)$$

The 2 in the denominator is included to account for the (close to) triangular shape of the abundance profiles of the proteins.

A second cost associated with the pinocytic vesicles, aside from the energy costs, is the area occupancy of under-construction vesicles on the plasma membrane. During the first stage of vesicle construction (Figure S1) the area blocked is assumed to increase linearly from zero to a disc shaped area with a radius equal to the vesicle radius. During the second stage this area is maintained. During the third stage the vesicle has been released from the plasma membrane so there is no blocking. These considerations yield an average area occupancy of:

$$\overline{A_{occ}} = \pi {r_{ves}}^2 \left( \frac{1}{2} \frac{t_1}{t_{UC}} + \frac{t_2}{t_{UC}} \right) \qquad (S26)$$

Summing over all under-construction vesicles and converting the occupied area into the number of nutrient transporters that would fit in that area yields:

$$N_{occ} = \frac{\pi {r_{ves}}^2 \left( \frac{1}{2} \frac{t_1}{t_{UC}} + \frac{t_2}{t_{UC}} \right) f_p \frac{t_{UC}}{t_{pers}} N_{ves,0}}{A_{mp}} \qquad (S27)$$

Simplifying and accounting for cell growth (see "Correcting for cell growth" section) yields the average number of nutrient transporters that are excluded over the cell cycle:

$$\overline{N_{occ}} = \frac{\pi {r_{ves}}^2 \left( \frac{1}{2} \frac{t_1}{t_{pers}} + \frac{t_2}{t_{pers}} \right) f_p N_{ves,0}}{A_{mp} ln(2)} \qquad (S28)$$

**Obtaining vesicle number from pinocytosis investment**

The gain in cell division time depends on the number of cytoplasmic vesicles which, in turn, depends on the investment in pinocytosis. The absolute investment in pinocytosis is the product of the relative investment in pinocytosis, $C_{pino}$, and the cost (or budget) of the entire cell at the start of the cell cycle, $C_{cell,0}$. Relating this product to the vesicle costs yields:

$$C_{pino}C_{cell,0} = C_{ves,0} + C_{UC,0} + C_{prot,0}$$
$$= 4\pi\left(r_{ves} - \frac{h_{mem}}{2}\right)^2 C_{area}N_{ves,0} + \overline{A_{UC}}C_{area}\frac{t_{UC}}{t_{pers}}N_{ves,0}$$
$$+ \frac{N_{ves,0}C_{AA}}{2t_{pers}}\sum_{i}^{All} N_{pp,i}\, t_{prot,i}N_{AA,i} \qquad (S29)$$

Rewriting yields the number of cytoplasmic vesicles:

$$N_{ves,0} = \frac{C_{pino}C_{cell,0}}{4\pi\left(r_{ves} - \frac{h_{mem}}{2}\right)^2 C_{area} + \overline{A_{UC}}C_{area}\frac{t_{UC}}{t_{pers}} + \frac{C_{AA}}{2t_{pers}}\sum_{i}^{All} N_{pp,i}\, t_{prot,i}N_{AA,i}} \qquad (S30)$$

The cell budget is obtained using[5]:

$$C_{cell,0} = C_\alpha {V_{cell,0}}^{0.97} \qquad (S31)$$

Where $C_\alpha$ is an empirically determined constant.

**Correcting for cell growth**

Depending on the requirements we use the nutrient transporter number, or other quantities, either at the start of the cell cycle ($N_{trans,0}$) or as an average over the cell cycle ($\overline{N_{trans}}$). The average requires that the growth of the cell is accounted for. As the cell grows throughout the cell cycle more plasma membrane transporters, vesicles, etc. are added that contribute to producing biomass. The cell volume grows exponentially because the increase in biomass depends linearly on the amount of biomass already present. The plasma membrane area is assumed to increase linearly with cell volume over the cell cycle, because the cell elongates as it grows. This simplifies the models and prevents a sudden jump in plasma membrane area at cell division (which would be required for a big spherical cell to divide into two smaller spherical cells).

When a fixed fraction of the plasma membrane is devoted to nutrient transporters the number of nutrient transporters on a single cell grows exponentially as the cell cycle progresses:

$$N_{trans}(t) = N_{trans,0}e^{at} \qquad (S32)$$

Here, $N_{trans,0}$ is the number of nutrient transporters in the plasma membrane at the start of the cell cycle. After the cell division time, $t_d$:

$$2N_{trans,0} = N_{trans,0}e^{at_d} \qquad (S33)$$

Solving this yields:

$$a = \frac{ln(2)}{t_d} \qquad (S34)$$

The average number of nutrient transporters over the cell cycle is given by:

$$\overline{N_{trans}} = \frac{\int_0^{t_d} N_{trans,0}e^{\frac{ln(2)}{t_d}t}dt}{t_d} = \frac{N_{trans,0}}{t_d}\int_0^{t_d} e^{\frac{ln(2)}{t_d}t}dt \qquad (S35)$$

Solving the integral yields:

$$\int_0^{t_d} e^{\frac{ln(2)}{t_d}t} dt = \frac{t_d}{ln(2)}\left(e^{ln(2)} - 1\right) = \frac{t_d}{ln(2)} \qquad (S36)$$

Plugging this back in yields:

$$\overline{N_{trans}} = \frac{N_{trans,0}}{t_d}\frac{t_d}{ln(2)} = \frac{N_{trans,0}}{ln(2)} \qquad (S37)$$

The same method can be used to obtain a growth corrected vesicle number.

**Solving the pinocytosis model**

To obtain the fitness, the Equations S13-S19, S22, and S30-S31 are plugged into Equation S11, leading to an expression that is fully determined by the parameter values and the variable, $C_{pino}$.

**Adjusting the pinocytosis model for larger vesicles**

The costs of vesicle construction have been experimentally determined for small vesicles only. The pinocytosis model can be applied to larger vesicles if we assume (1) that the geometry of the vesicle construction remains the same, and (2) that the cost of the proteins required to produce the vesicle scales linearly with under-construction vesicle surface area. The protein cost of vesicle production is modified to:

$$C_{prot,0} = \frac{A_{pillar,large}}{A_{pillar}}\frac{N_{ves,0}C_{AA}}{2t_{pers}}\sum_{i}^{All} N_{pp,i}\, t_{prot,i} N_{AA,i} \qquad (S38)$$

Where $A_{pillar,large}$ is the surface area of the large under-construction vesicles at their most extended state and $A_{pillar}$ is the equivalent for the small vesicles. To account for the larger cost of the membrane, $r_{ves}$ and $l$ are replaced by $r_{ves.large}$ and $l_{large}$.

**Number of nutrient transporters per vesicle in the pinocytosis model**

The number of nutrient transporters per vesicle is set by:

$$N_{trans,ves} = \frac{f_p(1 - f_{other})}{A_{mp}} 4\pi\left(r_{ves} - \frac{h_{mem}}{2}\right)^2 \qquad (S39)$$

Here, $f_p$ is the fraction of membrane surface area occupied by protein, $f_{other}$ is the fraction of membrane protein area that is occupied by membrane proteins other than nutrient transporters, $A_{mp}$ is the area on the membrane occupied by a single membrane protein, $r_{ves}$ is the vesicle radius, and $h_{mem}$ is the thickness of the membrane. The second part of the equation, $4\pi\left(r_{ves} - \frac{h_{mem}}{2}\right)^2$, is the area of the vesicle in the middle of the membrane, between the two leaflets of the lipid bilayer. A vesicle with a radius of 0.025 µm contains 25 nutrient transporters, whereas a vesicle with a radius of 0.1 µm contains 478 nutrient transporters.

### Proto-endoplasmic reticulum model

The proto-endoplasmic reticulum model compares two cellular states. An ancestral state that has a simple spherical cell lacking any internal membranes, with both nutrient transporters and Sec translocases localized to the plasma membrane. The Sec translocases insert integral membrane proteins with the assistance of ribosomes. In the derived state, both processes also exist on the plasma membrane, but in addition vesicles, that concentrate Sec translocases on their surface, bud from the plasma membrane. These vesicles serve as a primitive endoplasmic reticulum by inserting membrane proteins into their membranes and delivering them to the plasma membrane upon fusion. Lipids are assumed to insert into the vesicle membranes in parallel with membrane protein insertion so that membrane protein concentration remains constant. Lipid production is not treated explicitly, but in each parameter value combination used there is space on the vesicle membrane for additional enzymes.

The fitness of the derived state is calculated with respect to the ancestral state using Equation S11 (but using $C_{ER}$ and $t_{d,ER}$). This requires the determination of the cell division time of the ancestral state and the derived state.

#### Proto-endoplasmic reticulum—cell division time of the ancestral state

The cell division time of the ancestral state is calculated as in the pinocytosis model (Equation S13) but explicitly accounting for the extra membrane area that is used by Sec translocases. The average number of nutrient transporters is then given by:

$$\overline{N_{trans}} = \frac{4\pi\left(\frac{3V_{cell,0}}{4\pi}\right)^{\frac{2}{3}} f_p \left(1 - f_{other} - \frac{t_{ins}}{t_{d,anc}} ln(2)\right)}{A_{mp} ln(2)} \qquad (S40)$$

Here, $t_{ins}$ is the insertion time of a single membrane protein by a Sec translocase. The number of Sec translocases required depends on the cell division time because the longer an individual Sec translocon can insert membrane proteins the more it can insert and the fewer Sec translocons are required. Combining Equations S13 and S40 yields an equation that can be solved for $t_{d,anc}$.

#### Proto-endoplasmic reticulum—cell division time of the derived state

Calculating the cell division time of the derived state is different from that of the ancestral state in two respects. (1) Sec translocases are removed from the plasma membrane and replaced by nutrient transporters, increasing the number of nutrient transporters. (2) The vesicles used to remove the Sec translocases from the plasma membrane themselves occupy the plasma membrane, decreasing the number of nutrient transporters. Accounting for these two effects yields:

$$t_{d,ER} = \frac{N_{nutV} V_{cell,0}}{k_{cat} \frac{[nut]}{K_M + [nut]} \left(\overline{N_{trans}} - \left(\overline{N_{SecPM}} - \overline{N_{Sec,anc}}\right) - \overline{N_{occ}}\right)} \qquad (S41)$$

Here, $\overline{N_{trans}}$ is the average number of transporters present in the plasma membrane of the ancestral state (Equation S40), $\overline{N_{SecPM}}$ is the average number of Sec translocons present in the plasma membrane of the derived state, $\overline{N_{Sec,anc}}$ is the average number of Sec translocons present in the plasma membrane of the ancestral state. The combination $\left(\overline{N_{SecPM}} - \overline{N_{Sec,anc}}\right)$ yields a negative number because the vesicles reduce the number of Sec translocases present on the plasma membrane. Combined with the minus sign this yields an increase in the number of nutrient transporters in the plasma membrane. $\overline{N_{occ}}$ is the average number of nutrient transporters that is removed from the plasma membrane by the presence of under-construction vesicles. The membrane area occupancy of the nutrient transporters and Sec translocases is assumed to be the same, $10^{-5}$ µm$^3$.

$\overline{N_{trans}}$ is known from calculating the ancestral cell division time, leaving $\overline{N_{SecPM}}$, $\overline{N_{Sec,anc}}$, and $\overline{N_{occ}}$ as unknowns. All three can be calculated from known parameters, except that $\overline{N_{SecPM}}$ and $\overline{N_{occ}}$ are dependent also on the concentration of Sec translocon in the plasma membrane, $[Sec]_{PM}$. So, a fully assembled Equation S41 will contain two unknowns: $t_{d,ER}$ and $[Sec]_{PM}$. To solve Equation S41, it is combined with another equation that also depends on both $t_{d,ER}$ and $[Sec]_{PM}$, see below.

The number of Sec translocases on the plasma membrane in the derived state can be restated in terms of a concentration as:

$$\overline{N_{SecPM}} = \frac{{A_{pm,0}}^{*}[Sec]_{PM}}{ln(2)} \qquad (S42)$$

${A_{pm,0}}^{*}$ is the surface area of the plasma membrane immediately after cell division, but with the area that is blocked by the under-construction vesicles subtracted. ${A_{pm,0}}^{*}$ is obtained by multiplying the average area occupancy of a single vesicle, $\overline{A_{occ}}$ (Equation S26), by the number of under-construction vesicles, and subtracting that from the plasma membrane area:

$$\begin{aligned} {A_{pm,0}}^{*} &= A_{pm,0} - \pi {r_{ves}}^{2}\left(\frac{1}{2}\frac{t_1}{t_{UC}} + \frac{t_2}{t_{UC}}\right)\frac{t_{UC}}{t_{pers}}N_{ves,0} \\ &= A_{pm,0} - \pi {r_{ves}}^{2}\left(\frac{1}{2}\frac{t_1}{t_{pers}} + \frac{t_2}{t_{pers}}\right)N_{ves,0} \qquad (S43) \end{aligned}$$

Here, $\frac{t_{UC}}{t_{pers}}N_{ves,0}$ is the number of under-construction vesicles at the start of the cell cycle. $N_{ves,0}$ is the number of vesicles at the start of the cell cycle and is a function of the relative investment in the proto-endoplasmic reticulum, $C_{ER}$.

The average number of Sec translocons present in the plasma membrane of the ancestral state is given by:

$$\overline{N_{Sec,anc}} = \frac{A_{pm,0}f_p}{A_{mp}}\frac{t_{ins}}{t_{d,anc}} \qquad (S44)$$

With $\frac{A_{pm,0}f_p}{A_{mp}}$ being the total number of membrane proteins present in the new cell, and $\frac{t_{ins}}{t_{d,anc}}$ being the inverse of the number of membrane proteins that can be inserted into the membrane by a single Sec translocon in the ancestral state. The surface area of the plasma membrane, $A_{pm,0}$, is calculated from the cell volume, assuming that the cell is spherical.

The average number of nutrient transporters removed from the plasma membrane by the presence of vesicles, $\overline{N_{occ}}$, is like that of the pinocytosis model and given by Equation S28.

The number of vesicles present in the cell at the start of the cell cycle, $N_{ves,0}$, is calculated from the relative investment in the proto-ER:

$$N_{ves,0} = \frac{C_{ER}C_{cell,0}}{\overline{A_{ves}}C_{area} + \overline{A_{UC}}C_{area}\frac{t_{UC}}{t_{pers}} + \frac{C_{AA}}{2t_{pers}}\sum_i^{All} N_{pp,i}\, t_{prot,i}N_{AA,i}} \qquad (S45)$$

This differs from the pinocytosis model (Equation S30) in using an average vesicle area, $\overline{A_{ves}}$, because in the proto-endoplasmic reticulum model the vesicles grow as membrane proteins are added by the resident Sec translocases. The amount of membrane area added per unit time depends on the number of membrane proteins being added per unit time, which in turn is a function of the number of Sec translocases being present on the vesicle. The area of the vesicle as a function of time is:

$$A_{ves}(t) = \frac{A_{mp}}{f_p}N_{mpves}(t) \qquad (S46)$$

With

$$\begin{aligned} N_{mpves}(t^*) &= N_{mpves}(0) + \frac{1}{t_{ins}}\int_0^{t^*} N_{secves}(t)dt \\ &= N_{mpves}(0) + \frac{1}{t_{ins}}\int_0^{t^*} N_{secves}(0)e^{\frac{ln(2)}{t_{d,ER}}t}dt \qquad (S47) \end{aligned}$$

Here, $N_{mpves}(t)$ and $N_{secves}(t)$ are the number of membrane proteins and Sec translocases on the vesicle as a function of time. Sec translocases autocatalytically insert other Sec translocases and therefore grow in number exponentially (if the rest of the cell also grows exponentially). The rate constant of this exponential growth depends on the cell division time, and the fact that the cell doubles its contents over this period, hence the $\frac{ln(2)}{t_{d,ER}}$ (see Equation S32-S34). $N_{mpves}(0)$ is the number of membrane proteins in the vesicle just after vesicle production and is given by:

$$N_{mpves}(0) = \frac{f_pA_{ves,0}}{A_{mp}} \qquad (S48)$$

$N_{secves}(0)$ is the number of Sec translocases in the vesicle just after vesicle production and is given by:

$$N_{secves}(0) = \alpha[Sec]_{PM}A_{ves,0} \qquad (S49)$$

Here, $\alpha$ is a concentration factor for Sec translocases, from the plasma membrane into the vesicle. Substituting Equations S48 and S49 into Equation S47 and solving the integral yields:

$$N_{mpves}(t^*) = \frac{f_pA_{ves,0}}{A_{mp}} + \frac{\alpha[Sec]_{PM}A_{ves,0}}{t_{ins}}\frac{t_{d,ER}}{ln(2)}\left(e^{\frac{ln(2)}{t_{d,ER}}t^*} - 1\right) \qquad (S50)$$

Now the vesicle surface area becomes:

$$A_{ves}(t^*) = A_{ves,0} + \frac{A_{mp}}{f_p}\frac{\alpha[Sec]_{PM}A_{ves,0}}{t_{ins}}\frac{t_{d,ER}}{ln(2)}\left(e^{\frac{ln(2)}{t_{d,ER}}t^*} - 1\right) \qquad (S51)$$

When $\frac{ln(2)}{t_{d,ER}}t^* \ll 1$, the exponential function can be approximated as $e^x \approx x + 1$. This will be the case because $t^* = t_{pers}$, the persistence time of the fully formed vesicle in the cytoplasm, and the value of $t_{pers}$ will be kept to 0.1x of the cell division time, or smaller. This yields a vesicle surface area:

$$A_{ves}(t^*) = A_{ves,0} + \frac{A_{mp}}{f_p}\frac{\alpha[Sec]_{PM}A_{ves,0}}{t_{ins}}t^* \qquad (S52)$$

This means that over the lifetime of the vesicle, the Sec translocon amount on the vesicle effectively doesn't change. To obtain the average vesicle surface area Equation S52 needs to be integrated:

$$\overline{A_{ves}} = \frac{\int_0^{t_{pers}} A_{ves}(t)dt}{t_{pers}} \qquad (S53)$$

Yielding:

$$\overline{A_{ves}} = A_{ves,0} + \frac{1}{2}\frac{A_{mp}}{f_p}\frac{\alpha[Sec]_{PM}A_{ves,0}}{t_{ins}}t_{pers} \qquad (S54)$$

The vesicle surface area immediately after vesicle construction is given by:

$$A_{ves,0} = 4\pi\left(r_{ves} - \frac{h_{mem}}{2}\right)^2 \qquad (S55)$$

Here, $r_{ves}$ is the radius of the vesicle to the outer edge of the vesicle membrane, and $h_{mem}$ is the thickness of the membrane. $A_{ves,0}$ is the vesicle surface area in the middle of the membrane, to accurately account for membrane cost for small vesicles.

$\overline{N_{SecPM}}$, $\overline{N_{Sec,anc}}$, and $\overline{N_{occ}}$ can now be substituted in Equation S41, together with $\overline{N_{trans}}$, and an equation with two unknowns is obtained, $t_{d,ER}$ and $[Sec]_{PM}$.

*Second equation linking $t_{d,ER}$ and $[Sec]_{PM}$*

A second equation that also has $t_{d,ER}$ and $[Sec]_{PM}$ as unknowns can be established by equating the number of Sec translocases required to insert all the membrane proteins of a new cell, to the number of Sec translocases available on the plasma membrane and vesicles:

$$N_{Sec,needed,0} = N_{Sec,available,0} \qquad (S56)$$

Where the needed and available number of Sec translocases are compared at the start of the cell cycle. The number of Sec translocases that is needed depends on the total membrane area of a newly produced cell:

$$N_{Sec,needed,0} = \frac{A_{mem,0}f_p}{A_{mp}}\frac{t_{ins}}{t_{d,ER}}ln(2) \qquad (S57)$$

Here, $\frac{A_{mem,0}f_p}{A_{mp}}$ is the number of membrane proteins present in the cell (it is assumed that all membrane proteins have the same membrane area occupancy), $\frac{t_{ins}}{t_{d,ER}}$ is the inverse of the number of membrane proteins that can be inserted into the membrane by a single Sec translocon, and the factor $ln(2)$ is included to convert an average Sec translocon number needed to the number needed at the start of the cell cycle (compare with Equation S37). $N_{Sec,needed,0}$ only counts active Sec translocases.

$A_{mem,0}$ is the surface area of all the membrane in the cell at the start of the cell cycle and is given by:

$$A_{mem,0} = A_{pm,0}{}^{*} + A_{UCall,0} + A_{vesall,0} \qquad (S58)$$

$A_{pm,0}{}^{*}$ is the area of the plasma membrane with the area occupied by under-construction vesicles subtracted (Equation S43). $A_{UCall,0}$ is the combined surface area of all under-construction vesicles at the start of the cell division time. This is given by:

$$A_{UCall,0} = \overline{A_{UC}} \frac{t_{UC}}{t_{pers}} N_{ves,0} \qquad (S59)$$

Here, $\overline{A_{UC}}$ is the average surface area of a single under-construction vesicle and is given by Equation S22. The second part, $\frac{t_{UC}}{t_{pers}} N_{ves,0}$, is the number of under-construction vesicles present at the start of the cell cycle. With $t_{UC}$ being the construction time, $t_{pers}$ being the persistence time of an active (membrane protein inserting) vesicle in the cytoplasm, and $N_{ves,0}$ being the number of active vesicles present at the start of the cell cycle (Equation S45).

$A_{vesall,0}$ is the combined surface area of all active, membrane protein inserting, vesicles at the start of the cell cycle. It is calculated by:

$$A_{vesall,0} = \overline{A_{ves}} N_{ves,0} = \left( A_{ves,0} + \frac{1}{2} \frac{A_{mp}}{f_p} \frac{\alpha [Sec]_{PM} A_{ves,0}}{t_{ins}} t_{pers} \right) N_{ves,0} \qquad (S60)$$

Here, the average area of a single vesicle, $\overline{A_{ves}}$, is obtained from Equation S54, and the number of vesicles at the start of the cell cycle, $N_{ves,0}$, is obtained from Equation S45. Combining these results yields:

$$\begin{aligned} A_{mem,0} = A_{pm,0} & \\ & + \Bigg( \overline{A_{UC}} \frac{t_{UC}}{t_{pers}} + A_{ves,0} + \frac{1}{2} \frac{A_{mp}}{f_p} \frac{\alpha [Sec]_{PM} A_{ves,0}}{t_{ins}} t_{pers} \\ & - \pi r_{ves}{}^{2} \left( \frac{1}{2} \frac{t_1}{t_{pers}} + \frac{t_2}{t_{pers}} \right) \Bigg) N_{ves,0} \qquad (S61) \end{aligned}$$

Using Equation S45 for $N_{ves,0}$ and substituting $A_{mem,0}$ into Equation S57 yields the number of Sec translocases needed at the start of the cell cycle, $N_{Sec,needed,0}$.

This needs to be equated to the number of Sec translocases available at the start of the cell cycle, $N_{Sec,available,0}$, which is given by:

$$N_{Sec,available,0} = N_{SecPM,0} + N_{Secves,0} \qquad (S62)$$

$N_{SecPM,0}$ is the number of Sec translocases present on the plasma membrane at the start of the cell cycle and $N_{Secves,0}$ is the number of Sec translocases present in all active vesicles at the start of the cell cycle. There is no contribution from the under-construction vesicles because the Sec translocases present there are not active. The plasma membrane Sec translocase number is:

$$N_{SecPM,0} = [Sec]_{PM} A_{pm,0}{}^{*} \qquad (S63)$$

Here, $[Sec]_{PM}$ is the concentration of Sec translocon on the plasma membrane and $A_{pm,0}{}^{*}$ is the adjusted plasma membrane area (Equation S43). The total Sec translocon number in all active vesicles is given by:

$$N_{Secves,0} = \alpha[Sec]_{PM}A_{ves,0}N_{ves,0} \qquad (S64)$$

Here, $\alpha$ is the concentration factor for Sec translocases from the plasma membrane into vesicles and $A_{ves,0}$ is the surface area of a single vesicle immediately after it is constructed. The vesicle grows over time because membrane proteins (and implicitly the associated lipids) are inserted but the number of Sec translocons in the vesicles stays constant so to calculate the number of Sec translocases the amount at the start, $\alpha[Sec]_{PM}A_{ves,0}$, suffices. Combining Equations S63 and S64 yields:

$$N_{Sec,available,0} = [Sec]_{PM}A_{pm,0}{}^{*} + \alpha[Sec]_{PM}A_{ves,0}N_{ves,0} \qquad (S65)$$

Applying this result together with the $N_{Sec,needed,0}$ from Equation S57 to Equation S56, a new expression is obtained that depends on both $t_{d,ER}$ and $[Sec]_{PM}$.

Equation S41 can be substituted into Equation S56 so that a relation is obtained that depends only on $[Sec]_{PM}$. The resultant value of $[Sec]_{PM}$ can be plugged back into Equation S41 to obtain $t_{d,ER}$. Finally, this can be combined with $t_{d,anc}$ to find the fitness from Equation S11 (but using $C_{ER}$ and $t_{d,ER}$).

**Proto-endoplasmic reticulum—maximum attainable fitness**

A maximum attainable fitness can be calculated under two assumptions: (1) all Sec translocases are removed from the plasma membrane and replaced by nutrient transporters, and (2) this comes at no energetic cost or loss of plasma membrane area. The maximal fitness can be obtained by simplifying Equation S1:

$$Maximal\ fitness = e^{\ln(2)\left(1-\frac{t_d{}^{*}}{t_{d,anc}}\right)} \qquad (S66)$$

The cell division time of the ancestor is obtained from Equations S13 and S40, and simplified:

$$t_{d,anc} = \frac{\lambda}{\overline{N_{trans}}} \qquad (S67)$$

The cell division time of the derived state is:

$$t_d{}^{*} = \frac{\lambda}{\overline{N_{trans}} + \overline{N_{Sec,anc}}} \qquad (S68)$$

$\overline{N_{trans}}$ is the average number of nutrient transporters in the plasma membrane of the ancestor and $\overline{N_{Sec,anc}}$ is the average number of Sec translocases in the plasma membrane of the ancestor. The ratio of cell division times then becomes:

$$\frac{t_d{}^{*}}{t_{d,anc}} = \frac{\lambda/\left(\overline{N_{trans}} + \overline{N_{Sec,anc}}\right)}{\lambda/\overline{N_{trans}}} = \frac{\overline{N_{trans}}}{\overline{N_{trans}} + \overline{N_{Sec,anc}}} \qquad (S69)$$

$\overline{N_{trans}}$ and $\overline{N_{Sec,anc}}$ are given by Equations S40 and S44. Substituting and simplifying yields:

$$\frac{t_d{}^{*}}{t_{d,anc}} = \frac{1 - f_{other} - \frac{t_{ins}}{t_{d,anc}}ln(2)}{1 - f_{other}} \qquad (S70)$$

Here, $f_{other}$ is the fraction of the plasma membrane area not devoted to nutrient transport and membrane protein insertion, $t_{ins}$ is the insertion time of a single membrane protein, and $t_{d,anc}$ is the ancestral cell division time. After substitution the maximal fitness is given by:

$$Maximal\ fitness = e^{\ln(2)\left(1-\frac{1-f_{other}-\frac{t_{ins}}{t_{d,anc}}ln(2)}{1-f_{other}}\right)} \qquad (S71)$$

The minimal cell division time associated with the maximal fitness is given by:

$$t_d{}^* = t_{d,anc}\left(1-\frac{ln(Maximal\ fitness)}{\ln(2)}\right) \qquad (S72)$$

**Calculating the vesicle Sec translocase limit imposed by ribosomes**

Membrane proteins are inserted into the membrane co-translationally, with the ribosomes docking on the Sec translocase. Ribosomes are considerably larger than Sec translocases and their steric interaction limits the number of Sec translocases that can function on a single vesicle. To estimate the number of ribosomes that will fit, it is assumed that each ribosome is a sphere with a 10 nm radius. The midplanes of the ribosomes, with an area occupancy of $\pi r_{ribo}{}^2$, sit on a surface 10 nm above the surface of the vesicle, this surface initially has an area of $4\pi(r_{ves}+r_{ribo})^2$. The initial maximal number of ribosomes is given as $\frac{4\pi(r_{ves}+r_{ribo})^2}{\pi r_{ribo}{}^2}$. After a single round of insertions has been completed the area of the vesicle has increased which allows more ribosomes to fit. This continues with the second round, etc. After each round of insertion, the maximal number of ribosomes is recalculated. The maximal number of ribosomes, and therefore Sec translocases, is artificially limited to a maximum of 100 so that there is space for other activities on the vesicle, such as lipid production and vesicle fusion machinery. The maximal number of membrane proteins on a vesicle at the start is 254, using a radius of 0.0225 µm (the middle of the bilayer in a 0.05 µm diameter vesicle), a membrane protein area fraction of 0.4, and a membrane protein area occupancy of $10^{-5}$ µm$^2$. The ribosome limit used in the plots is an average of the number of ribosomes that fit over the lifetime of the vesicle.

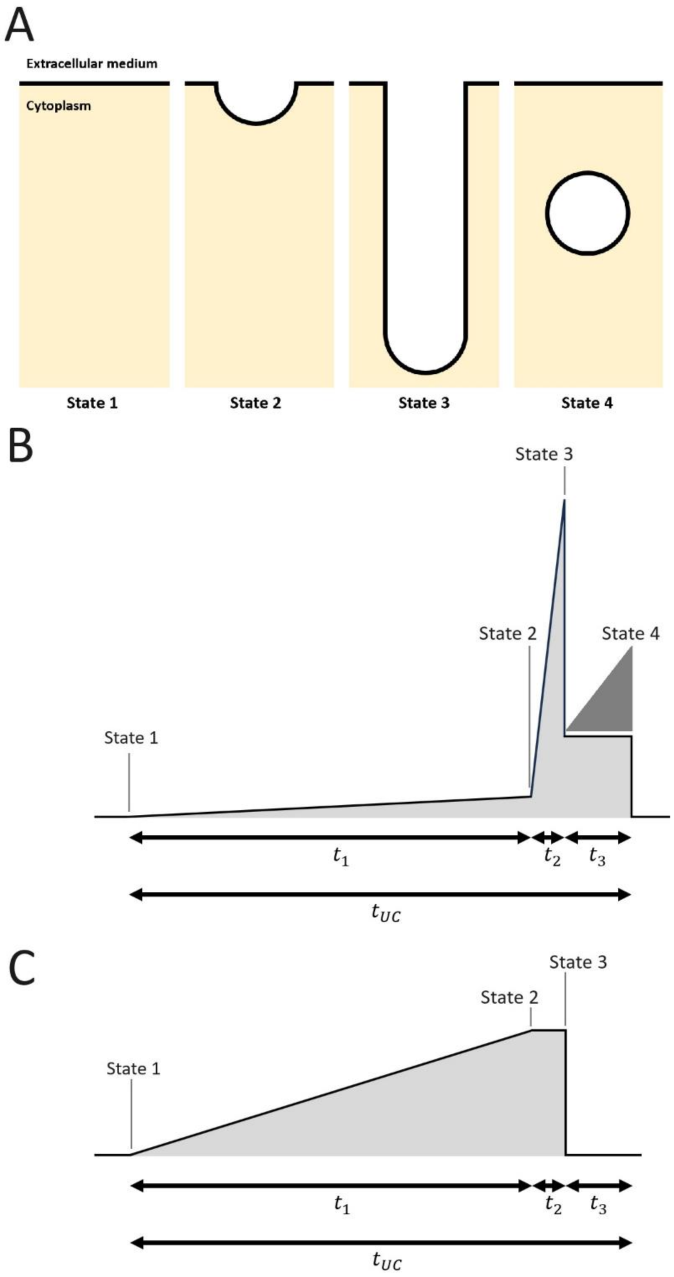


**Figure S1. Related to Figure 1 and 4. Schematic of the production of vesicles at the plasma membrane.**
(A) Different states of vesicle construction. State 1 is the starting point with a flat plasma membrane and without any vesicle constructing proteins being present. From state 1 onward, vesicle construction proteins associate with the plasma membrane and form a bulge. This culminates in the hemisphere of state 2. Starting after state 2 the membrane is pulled outward forming a cylinder topped with a hemisphere culminating in state 3. Immediately after state 3 the vesicle membrane is completed, and persists as state 4. During state 4 the vesicle is still inactive and is therefore still counted as an under-construction vesicle. Stage 1 from the text is the period between state 1 and 2, stage 2 is the period between state 2 and 3, and stage 3 is the period the vesicle is in state 4.
(B) Change in the area of the under-construction vesicle (minus the area missing from the plasma membrane) over time. The times $t_1$, $t_2$, $t_3$, and $t_{UC}$ are model parameters.
(C) Change in the area of the plasma membrane that is blocked by the under-construction vesicle over time.

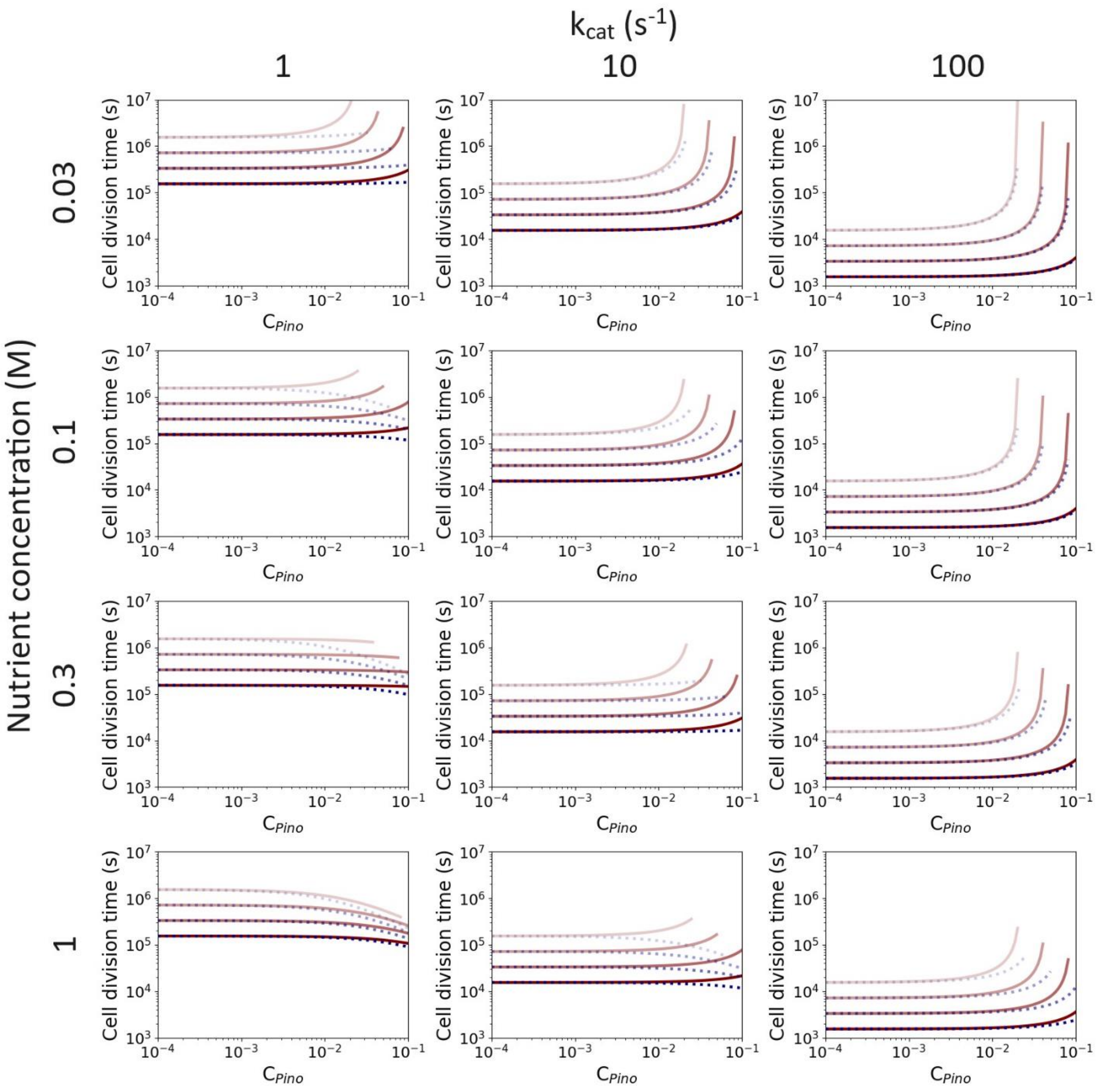


**Figure S2. Related to Figure 2. Cell division time of the derived state in the pinocytosis model.**
Red solid lines: vesicle radius 0.025 µm. Blue dotted lines: vesicle radius 0.1 µm. The cell volume varies from 10, $10^2$, $10^3$, to $10^4$ µm$^3$ with the darker lines (for both red and blue) being the smaller volumes. At some level of investment in pinocytosis, the under-construction vesicles block the nutrient transporter area on the plasma membrane completely. Here the model breaks down and the plot is cut off.

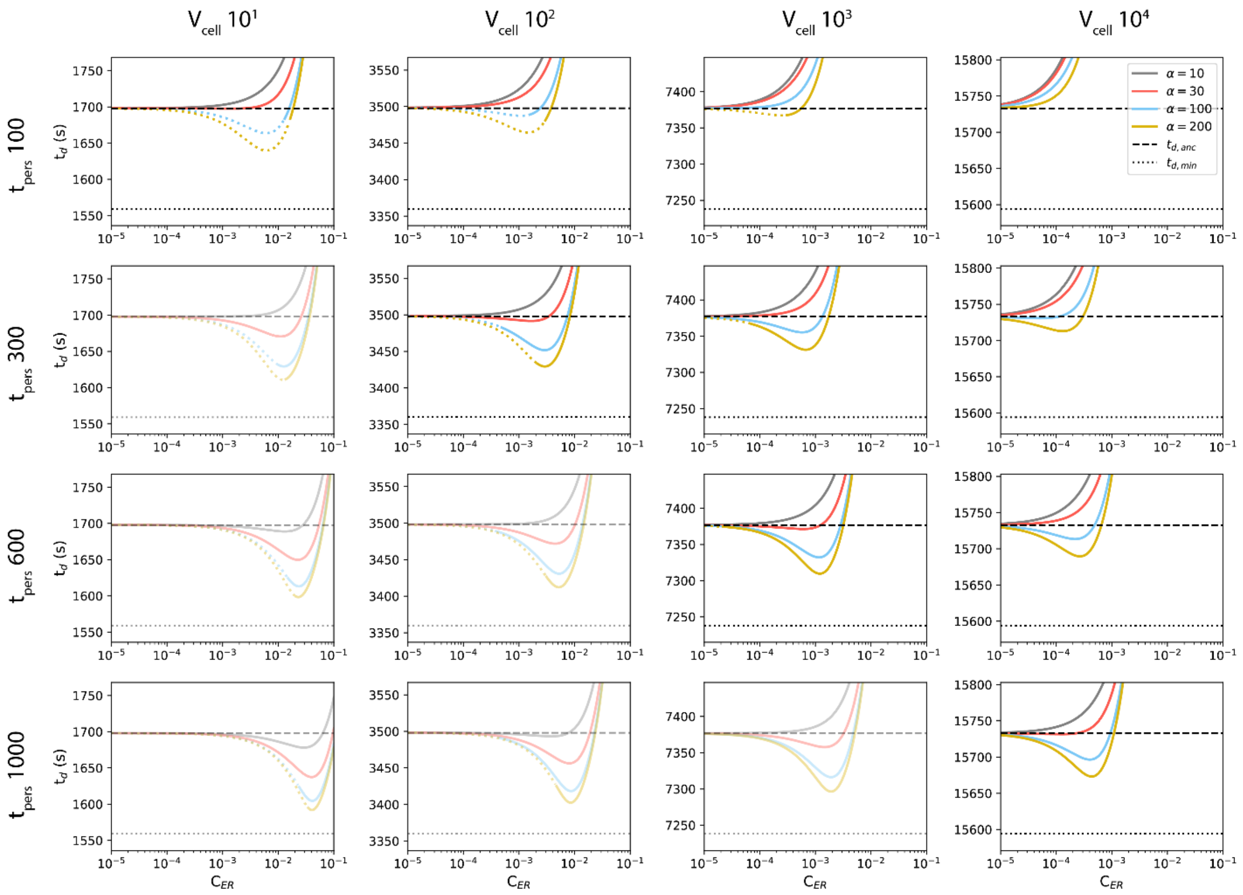


**Figure S3. Related to Figure 5. Cell division time of the derived state for a $t_{ins}$ of 20 s.**

Cell volumes ($V_{cell}$) in μm$^3$, vesicle persistence times ($t_{pers}$) in seconds. The parameter α shows the concentration factor for Sec translocases from plasma membrane into vesicles. Dotted colored lines show where the model breaks down due to ribosome packing on the vesicle surface. Solid colored lines show where the model works. Black dashed lines: the cell division time of the ancestral state. Black dotted lines: the minimal attainable cell division time. Faded plots indicate the parameter combinations for which $t_{pers}/t_d > 0.1$. To derive the model, it was assumed that $t_{pers}/t_d \ll 1$.

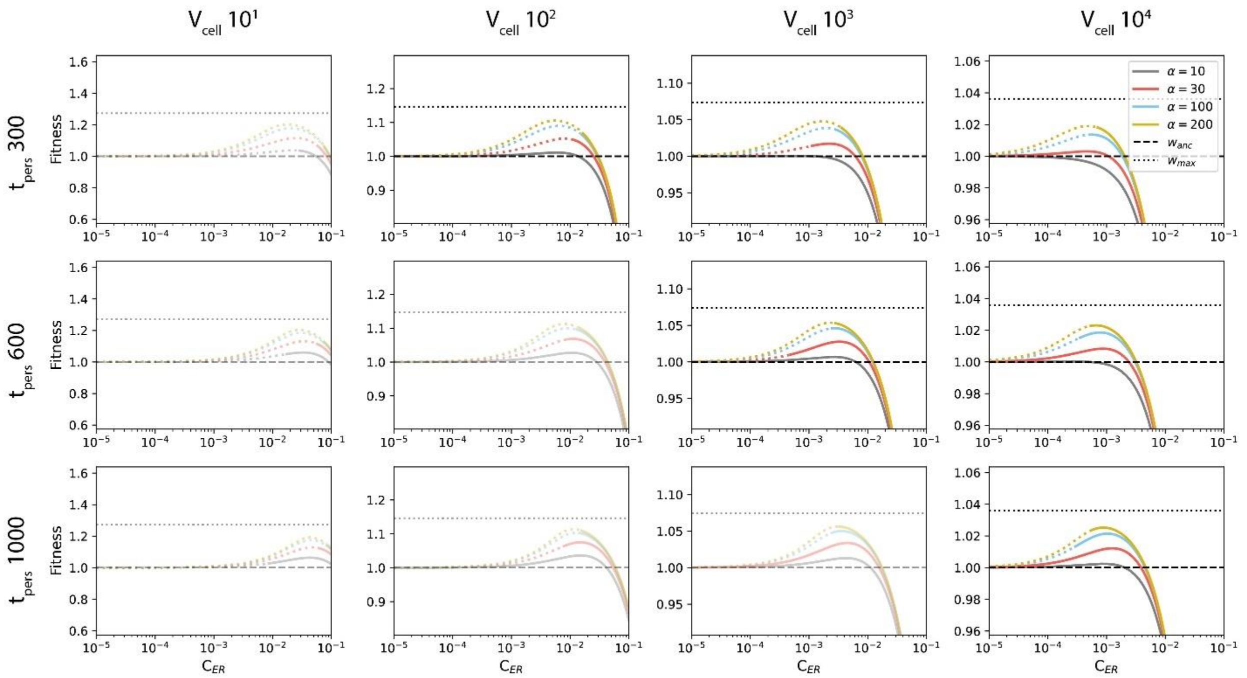


**Figure S4. Related to Figure 6A. Proto-endoplasmic reticulum fitness for a $t_{ins}$ of 120 s.**

The fitness of the derived state, containing a proto-endoplasmic reticulum, as a function of the relative cost of the proto-endoplasmic reticulum. The insertion time for individual membrane proteins ($t_{ins}$) is 120 s. Cell volumes ($V_{cell}$) in $\mu m^3$, vesicle persistence times ($t_{pers}$) in seconds. The parameter α shows the concentration factor for Sec translocases from plasma membrane into vesicles. Dotted colored lines show where the model breaks down due to ribosome packing on the vesicle surface. Solid colored lines show where the model works. Dashed black lines: the fitness of the ancestor, dotted black lines: the maximal attainable fitness. Faded plots indicate the parameter combinations for which $t_{pers}/t_d > 0.1$. To derive the model, it was assumed that $t_{pers}/t_d \ll 1$.

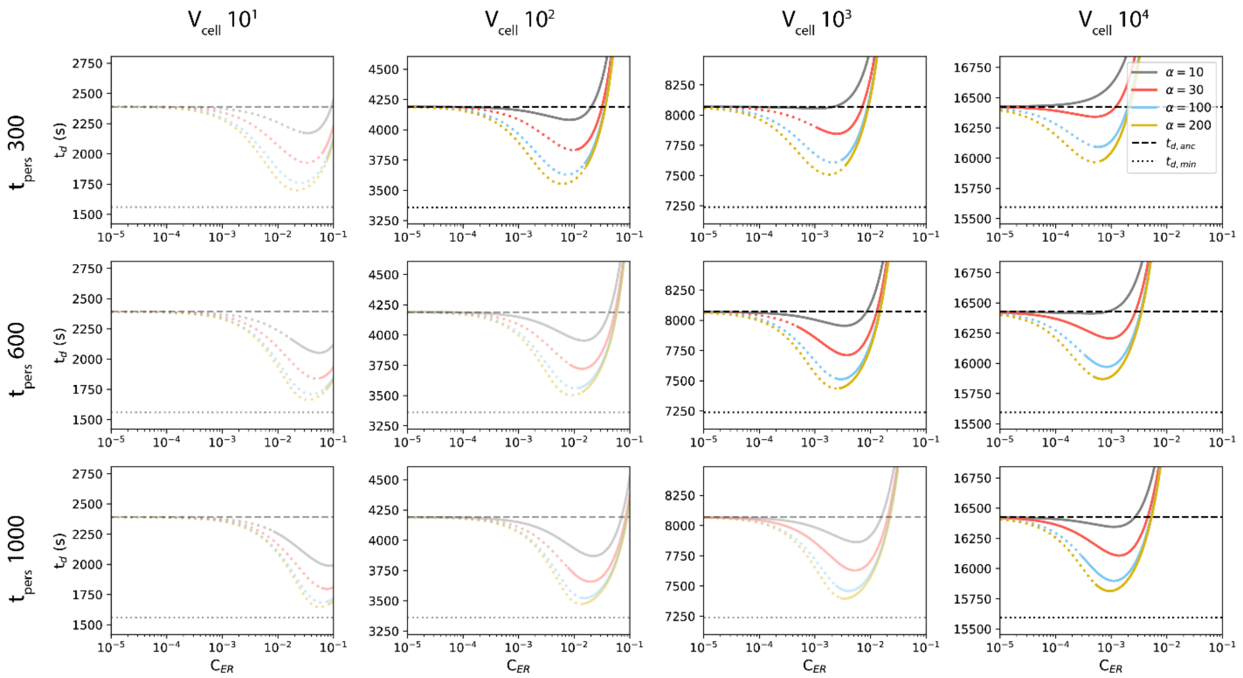


**Figure S5. Related to Figure 6A. Cell division time of the derived state for a $t_{ins}$ of 120 s.**
Cell volumes ($V_{cell}$) in µm³, vesicle persistence times ($t_{pers}$) in seconds. The parameter α shows the concentration factor for Sec translocases from plasma membrane into vesicles. Dotted colored lines show where the model breaks down due to ribosome packing on the vesicle surface. Solid colored lines show where the model works. Black dashed lines: the cell division time of the ancestral state. Black dotted lines: the minimal attainable cell division time. Faded plots indicate the parameter combinations for which $t_{pers}/t_d > 0.1$. To derive the model, it was assumed that $t_{pers}/t_d \ll 1$.

**Table S1. Related to Figure 2. Vesicle persistence times for small vesicles in the pinocytosis model (50 nm diameter).**

| | $k_{cat}$ = 1 $s^{-1}$ | $k_{cat}$ = 10 $s^{-1}$ | $k_{cat}$ = 100 $s^{-1}$ |
|---|---|---|---|
| **[Nut] = 0.03 M** | 23.8 | 2.38 | 0.238 |
| **[Nut] = 0.1 M** | 79.3 | 7.93 | 0.793 |
| **[Nut] = 0.3 M** | 238 | 23.8 | 2.38 |
| **[Nut] = 1 M** | 793 | 79.3 | 7.93 |

**Table S2. Related to Figure 2. Vesicle persistence times for large vesicles in the pinocytosis model (200 nm diameter).**

| | $k_{cat}$ = 1 $s^{-1}$ | $k_{cat}$ = 10 $s^{-1}$ | $k_{cat}$ = 100 $s^{-1}$ |
|---|---|---|---|
| **[Nut] = 0.03 M** | 136 | 13.6 | 1.36 |
| **[Nut] = 0.1 M** | 453 | 45.3 | 4.53 |
| **[Nut] = 0.3 M** | 1358 | 135.8 | 13.58 |
| **[Nut] = 1 M** | 4526 | 452.6 | 45.26 |

**Table S3. Related to Figure 1 and 4. Quantification of proteins used in vesicle construction.**
Protein names, peak copy number, and time present are for Schizosaccharomyces pombe[4]. Protein lengths were obtained from KEGG (T00076, SPAC26A3.05). App1p was not found in KEGG, so the protein length was extracted from Pombase (SPBC29B5.04c). ARPC5 wasn't found in KEGG, instead Arc5 was used for protein length.

| Protein name | Protein length (amino acids) | Peak copy number in vesicle | Time present in vesicle (s) |
|---|---|---|---|
| Chc1p | 1666 | 40 | 110 |
| Clc1p | 229 | 40 | 115 |
| End4p | 1102 | 160 | 41 |
| Pan1p | 1794 | 260 | 41 |
| Wsp1p | 574 | 230 | 12 |
| Vrp1p | 309 | 140 | 11 |
| Myo1p | 1217 | 400 | 14 |
| Arp2 | 390 | 320 | 26 |
| Arp3 | 427 | 320 | 26 |
| ARPC5 | 152 | 320 | 26 |
| Fim1p | 614 | 910 | 22 |
| Acp2p | 268 | 230 | 20 |
| App1p | 605 | 150 | 15 |
| Crn1p | 601 | 490 | 21 |
| Twf1p | 328 | 210 | 17 |
| Actin | 375 | 7500 | 19 |

1. Lynch, M., and Marinov, G.K. (2017). Membranes, energetics, and evolution across the prokaryote-eukaryote divide. Elife *6*, e20437.

2. Schavemaker, P.E., and Lynch, M. (2022). Flagellar energy costs across the tree of life. Elife *11*, e77266.

3. Lindén, M., Sens, P., and Phillips, R. (2012). Entropic Tension in Crowded Membranes. PLoS Comput Biol *8*, 1–10. 10.1371/journal.pcbi.1002431.

4. Sirotkin, V., Berro, J., Macmillan, K., Zhao, L., and Pollard, T.D. (2010). Quantitative analysis of the mechanism of endocytic actin patch assembly and disassembly in fission yeast. Mol Biol Cell *21*, 2894–2904.

5. Lynch, M., and Marinov, G.K. (2015). The bioenergetic costs of a gene. Proceedings of the National Academy of Sciences *112*, 15690–15695.
